# Microglial inflammatory activation paradoxically improves glucose tolerance during diet-induced obesity

**DOI:** 10.1101/2022.04.19.488819

**Authors:** J.D. Douglass, M. Valdearcos, K.M. Ness, A. Wyse-Jackson, M.D. Dorfman, J. M. Frey, A. Niraula, J. Banerjee, R.D. Fasnacht, M. Robblee, S.K. Koliwad, J.P. Thaler

## Abstract

Hypothalamic gliosis associated with high fat diet (HFD) feeding increases susceptibility to hyperphagia and weight gain, and is therefore presumed to promote obesity-associated consequences such as glucose intolerance as well. Nevertheless, the body weight-independent contribution of microglial activation to glucose regulation has not been determined. Here we show that reducing microglial NF-κB signaling via cell-specific IKKβ deletion exacerbates HFD-induced glucose intolerance and insulin resistance despite reducing body weight and adiposity. This effect was associated with reduced activity of hypothalamic glucose sensing neurons. Conversely, a genetic approach to increase microglial inflammatory activity improved glucose tolerance independently of diet in lean rodents. To avoid confounding effects due to chronic alterations to microglial signaling pathways from dietary or genetic interventions, we developed an inducible model of microglial activation using DREADD-based chemogenetics. Gq-coupled GPCR activation rapidly increased microglial calcium levels, cytokine gene expression, and morphological hallmarks of inflammatory activation. In both lean and obese rodents, chemogenetic microglial activation caused a marked improvement in glucose tolerance along with increased activation of hypothalamic glucose sensing neurons, effects abrogated by central blockade of TNFα signaling. Thus, while diet-induced microglial activation promotes weight gain, it may also serve an adaptive function—to prevent the deterioration of glucose tolerance associated with obesity, an important consideration for immune-modulating metabolic therapies.

## Introduction

In humans and rodents, obesity is associated with gliosis, the inflammatory activation of non-neuronal cells such as microglia (the resident macrophages of the CNS) (Ávalos et al., 2018; Berkseth et al., 2014; Douglass et al., 2017a; Schur et al., 2015; Thaler et al., 2012; Valdearcos et al., 2015, 2019). This gliosis occurs at the onset of high-fat diet (HFD) feeding and is largely restricted to brain regions such as the hypothalamus that contain neurocircuits critical to the regulation of body weight and glucose homeostasis (Berkseth et al., 2014; Douglass et al., 2017a; Gao et al., 2014; Schur et al., 2015; Thaler et al., 2012; Valdearcos et al., 2015, 2019). These observations led to the hypothesis that activated microglia promote the progression of diet-induced obesity (DIO) and insulin resistance (Ávalos et al., 2018; Berkseth et al., 2014; Douglass et al., 2017a; Schur et al., 2015; Thaler et al., 2012; Valdearcos et al., 2015, 2019). Indeed, there is now substantial evidence that microglia increase DIO susceptibility (Dorfman et al., 2017; Gao et al., 2017; Kim et al., 2019; Niraula et al., 2021; Valdearcos et al., 2014, 2017; Wang et al., 2020), but a weight-independent role for microglia in glucose homeostasis has not been established.

Microglia are long-lived CNS-resident immune cells that are intimately involved in both physiological and pathological processes within the brain (Colonna and Butovsky, 2017; Prinz et al., 2019, 2021; Salter and Stevens, 2017). Microglia regulate many homeostatic functions in the healthy brain, but their plasticity and sensitivity to environmental cues elicit rapid transformation to more reactive cellular phenotypes in disease states (Colonna and Butovsky, 2017; Douglass et al., 2017a; Prinz et al., 2019, 2021; Salter and Stevens, 2017; Valdearcos et al., 2015). This cellular activation can have beneficial or deleterious impacts on surrounding neurons and glia, resulting in attenuation or exacerbation of pathology depending on the context. In DIO, for example, HFD induction of NF-κB-mediated signaling in microglia promotes food intake, fat accumulation, and central resistance to satiety signals such as leptin (Valdearcos et al., 2014, 2017). These data together with more recent analyses of microglial mitochondrial (UCP2) (Kim et al., 2019), lipid (LPL) (Gao et al., 2017), circadian (Bmal1) (Wang et al., 2020), chemokine (CX3CR1) (Dorfman et al., 2017), and prostaglandin (Niraula et al., 2021) signaling support a primary role for microglial activation in obesity pathogenesis.

Glucose intolerance and insulin resistance typically arise as a consequence of obesity. Therefore, the studies of microglial alterations to body weight showed the expected parallel effects on glycemic parameters (Dorfman et al., 2017; Gao et al., 2017; Kim et al., 2019; Niraula et al., 2021; Wang et al., 2020). However, CNS regulation of energy balance and systemic glycemia occurs via distinct pathways (Myers et al., 2021), raising the untested possibility of direct body weight-independent effects of microglial activation on glucose homeostasis. Here, using cell-specific models combined with approaches to account for or eliminate weight differences between groups, we reveal an unexpected benefit of microglial activation to improve glucose tolerance through a TNFα-dependent mechanism.

## Results

### Microglia-specific deletion of IKKβ ameliorates HFD-induced weight gain without accompanying improvements in glucose tolerance

We previously demonstrated that microglial activation induced by HFD feeding increases food intake and weight gain in mice (Dorfman et al., 2017; Niraula et al., 2021; Valdearcos et al., 2014, 2017), but the impact on glucose homeostasis was not explored. To examine the impact of microglial inflammatory signaling on glycemia in established obesity, we returned to our previously published *Cx3cr1*^CreER/+^::*Ikbkb*^fl/fl^ mouse model (IKKβ-MGKO) (Valdearcos et al., 2017). In this inducible model, tamoxifen treatment causes deletion of IKKβ in both peripheral and CNS Cx3cr1+ myeloid populations, but peripheral cells turn over after 4 weeks with replacement by nonrecombined bone marrow-derived monocytes (Dumas et al., 2021; Goldmann et al., 2013; Parkhurst et al., 2013; Yona et al., 2013). In contrast, microglia are long-lived with minimal turnover resulting in persistent gene deletion for >8 months (data not shown; (Dumas et al., 2021; Goldmann et al., 2013; Parkhurst et al., 2013; Yona et al., 2013)). Consistent with our previous findings (Valdearcos et al., 2017), NF-κB pathway silencing in IKKβ-MGKO mice resulted in a greatly diminished cytokine response to inflammatory stimulation with lipopolysaccharide (LPS) in primary microglia (Figure 1A). 4 weeks after tamoxifen treatment, control (*Ikbkb*^fl/fl^, Ctl) and IKKβ-MGKO mice were exposed to *ad libitum* low-fat control diet (CD) or 60% HFD for 10 weeks. Despite the >10% lower body weight and fat mass in HFD-fed IKKβ-MGKO mice compared to HFD-fed controls (Figures 1B and 1C), glucose tolerance was relatively unaffected (Figures 1D and 1E), a result that contrasts with those seen in other models with a similar degree of protection against DIO in the context of reduced CNS inflammation (Douglass et al., 2017b; Kleinridders et al., 2009; Zhang et al., 2008, 2017). Though CD-fed mice of both genotypes show only a slight uptrend in glucose AUC with increasing fat mass likely due to the low variance in the distribution of adiposity (Figure 1F), the HFD-fed control mice show the expected association of glucose intolerance with obesity (Figure 1G; gray dots, R^2^=0.51, p=0.01). In contrast, no such correlation was observed in HFD-fed IKKβ-MGKO mice (Figure 1G; red dots, R^2^=0.01, p=0.75) with a large proportion of leaner animals showing substantial glucose intolerance (upper left portion of graph in Figure 1G). Overall, these results suggest that loss of microglial inflammatory signaling during HFD exposure has divergent effects on the central regulation of body weight and glucose homeostasis.

**Figure 1.**
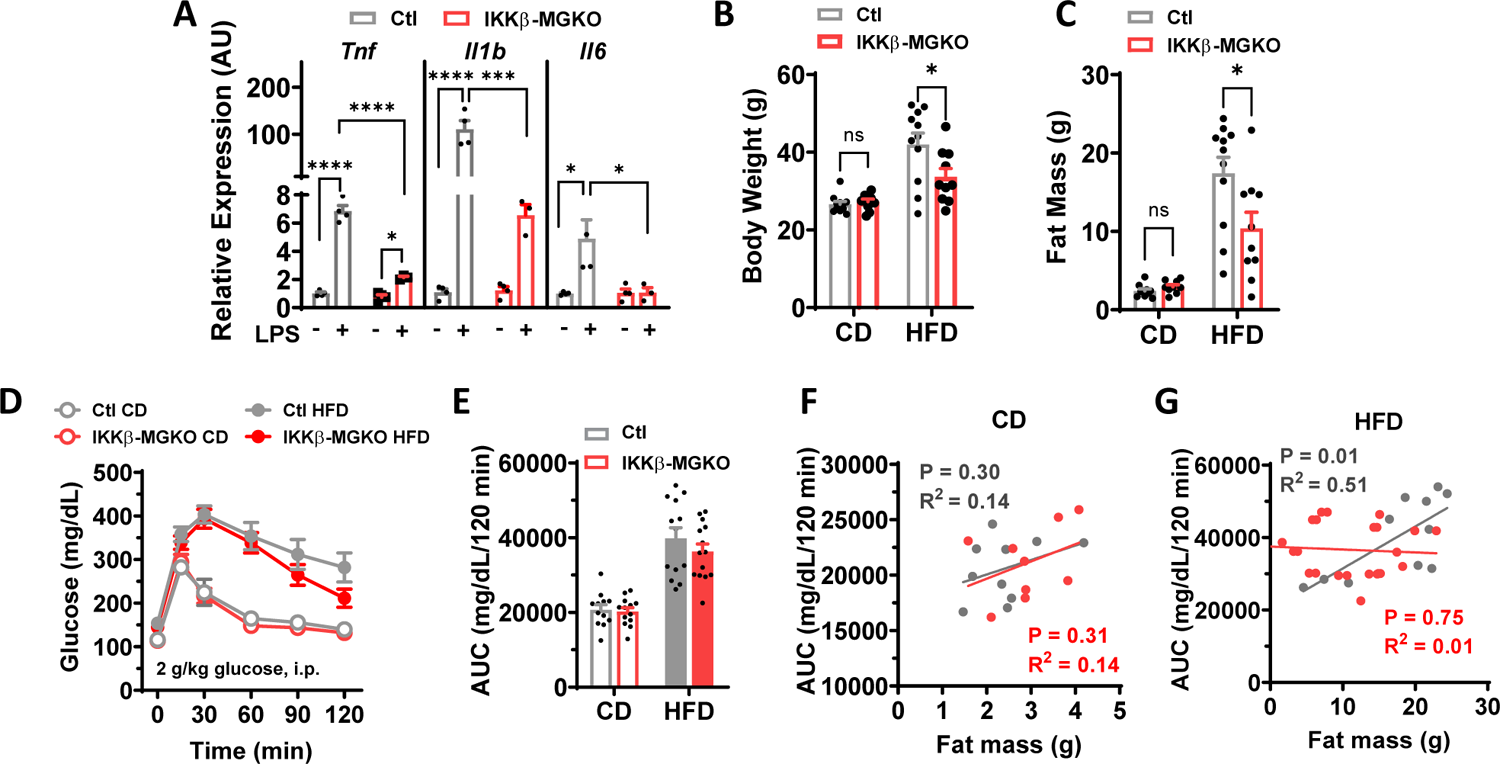
Loss of microglial IKKβ dampens inflammatory signaling and reduces HFD-induced weight gain but does not improve glucose tolerance. (A) Relative gene expression of inflammatory cytokines in primary microglia cultures from IKKβ-MGKO and control (Ctl) mice treated for 6 hours with 100 ng/ml lipopolysaccharide (LPS). N=3-4 wells/genotype/treatment. Two-way ANOVA with Bonferroni post-hoc test, * p < 0.05, *** p < 0.001, **** p <0.0001. (B) Body weight and (C) total fat mass after 10 weeks of *ad libitum* CD and HFD feeding. N=10-12/group. Two-way ANOVA with Bonferroni post-hoc test, * p < 0.05. (D) GTT (2 g/kg i.p.) in 4 hour fasted mice at 10 weeks of CD or HFD. N=10-12/group. (E) Total blood glucose area under the curve (AUC) for the GTT. (F-G) Linear regressions of total AUC versus 10 week weight gain in (F) CD-fed and (G) HFD-fed IKKβ-MGKO (red) and Ctl (gray) mice. All values are mean ± SEM.

### Loss of microglia NF-κB-mediated inflammation during HFD exposure exacerbates glucose intolerance and insulin resistance independently of body weight

Given the unexpected lack of improvement in glucose tolerance in the HFD-fed IKKβ-MGKO mice relative to controls, we wondered whether controlling for differences in body weight might actually reveal a <u>deterioration</u> of glycemic parameters in the knockout mice. Lower food intake is the primary cause of leanness in this model (Valdearcos et al., 2017). Accordingly, we generated weight-matched cohorts by pairing the HFD caloric consumption in control mice to the reduced intake of *ad libitum* (AL) HFD-fed IKKβ-MGKO mice. As predicted, this paradigm resulted in similarly matched body weights (Figure 2A) and adiposity (Figure 2B) between pair-fed controls and IKKβ-MGKO mice over 14 weeks of HFD consumption. Importantly, early dark cycle food intake and random blood glucose levels measured 4 hours after dark onset were similar between groups (Figures S1A and S1B), indicating that this mild pair feeding regimen did not induce the hyperphagia-fasting cycles common to many caloric restriction protocols. Nevertheless, to avoid any potential confounding from differing meal patterns, we provided *ad lib* HFD access to both groups for at least 2 days before assessing glucose homeostasis. At week 4 of HFD, IKKβ-MGKO mice displayed worse glucose tolerance than the weight-matched controls (Figures 2C and 2D), an effect that was sustained through the 12 weeks of HFD feeding (data not shown). Taken together these data indicate that microglial inflammatory activation is required to maintain normal glucose homeostasis despite promoting hyperphagia and increased adiposity under obesogenic dietary conditions.

**Figure 2.**
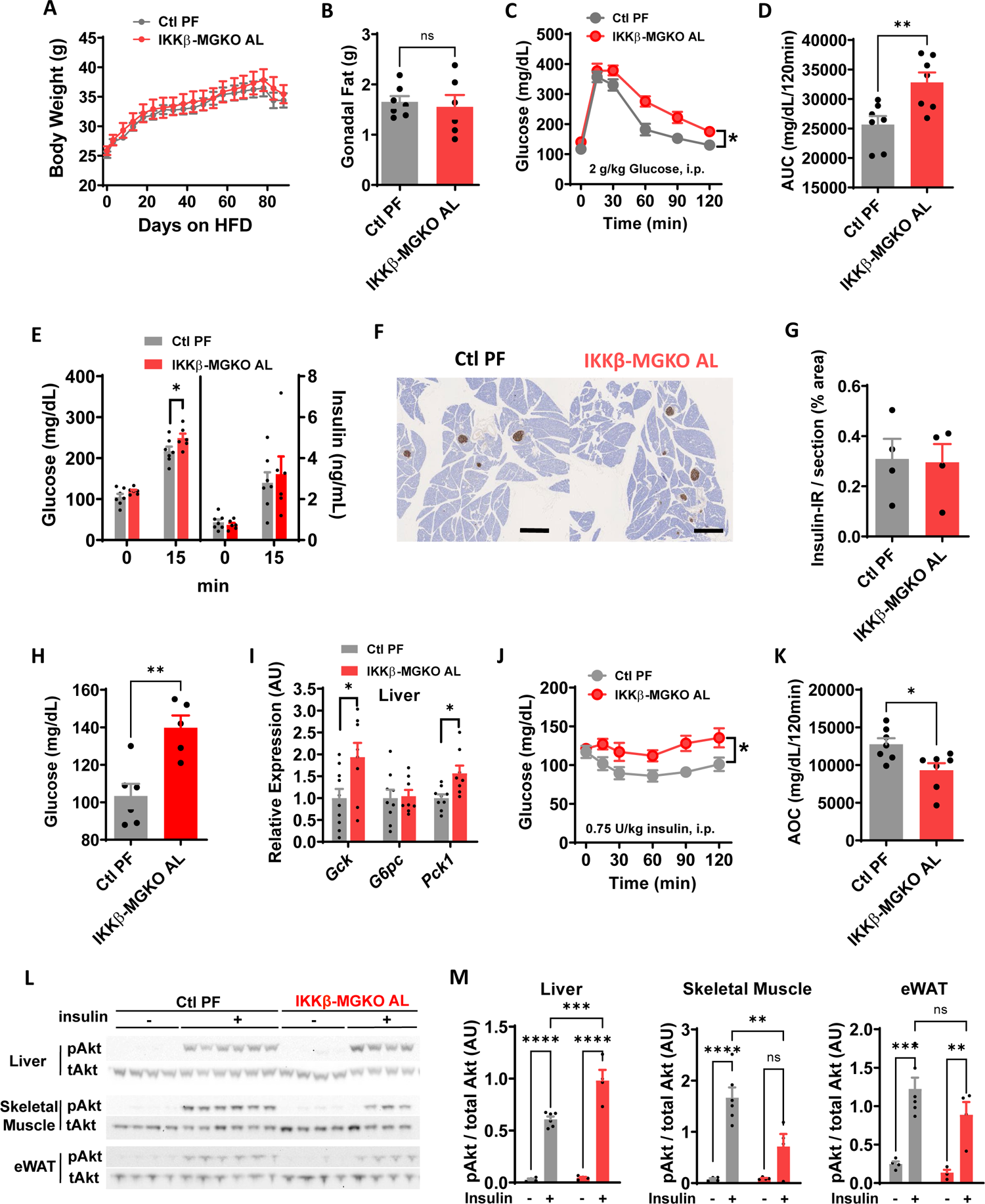
Microglial IKKβ deletion exacerbates HFD-induced glucose intolerance and insulin resistance in weight-matched mice. (A) Body weight of *ad libitum* HFD-fed IKKβ-MGKO (IKKβ-MGKO AL) and HFD pair-fed control mice (Ctl PF). N=7/group. (B) Gonadal fat pad weights after 14 weeks of HFD. (C) GTT (2 g/kg glucose, i.p.) and (D) total glucose AUC at 7 weeks of HFD. Two-way ANOVA and Student’s *t-*test, * p < 0.05, *** p < 0.001. (E) Glucose-stimulated insulin secretion (2 g/kg glucose by gavage). Student’s *t-*test, * p < 0.05 (F) Insulin (brown) immunostaining in fixed pancreatic sections of 10 week HFD-fed mice. Scale bar = 500 μm. (G) Quantification of insulin immunoreactivity. N=4/group. (H) 16-h fasted blood glucose. Student’s *t-*test, ** p < 0.01. (I) Liver mRNA levels of glucokinase (*Gck*), glucose-6-phosphatase (*G6pc*), and phosphoenolpyruvate carboxykinase 1 (*Pck1*) in overnight fasted mice after 14 weeks of HFD. N=8-10/group. Values normalized to Ctl PF expression. Student’s t-test, * p < 0.05. (J) ITT (0.75 U/kg insulin i.p.) and (K) Area-over-curve (AOC) at 10 weeks HFD. Two-way ANOVA and Student’s *t-*test, * p < 0.05 (L) pAkt (Ser473) and total Akt in tissue homogenates skeletal muscle, liver, and epididymal white adipose tissue (eWAT) from 16-h fasted mice given saline (-) or insulin (+) (5 U/kg, i.p., 15 minutes) after 14 weeks on HFD. (M) Densitometric quantification of tissues from L. N=4-6/group. Two-way ANOVA with Bonferroni post-hoc test, ** p < 0.01, *** p < 0.001, **** p < 0.0001. All values are mean ± SEM.

Next, we determined whether impairments in insulin secretion or action might underlie the glucose intolerance of IKKβ-MGKO mice. We performed an oral glucose challenge on IKKβ-MGKO and control mice and found that plasma insulin levels measured both in the fasted state and 15 minutes after glucose intake were similar between groups (Figure 2E). Moreover, immunohistochemical analysis of pancreatic islets after 10 weeks of HFD feeding revealed similar levels of pancreatic insulin content across genotypes (Figures 2F and 2G). These data together suggest that IKKβ-MGKO do not have deficient insulin production.

IKKβ-MGKO mice had elevated blood glucose levels after fasting overnight for 16 hours (Figure 2H), suggesting a potential impairment of insulin action at the liver. Analyses of hepatic mRNA in fasted mice showed transcriptional upregulation of the primary gluconeogenic enzymes phosphoenolpyruvate carboxykinase (*Pck1*) and glucokinase (*Gck*) with no change in glucose-6 phosphatase (G6pc) (Figure 2I), analogous to that observed in pre-diabetic animals with enhanced endogenous glucose production (Ferre et al., 2003; Torres et al., 2009). In addition, a pyruvate tolerance test revealed modestly elevated blood glucose in the IKKβ-MGKO mice relative to controls (Figures S1C and S1D).

Consistent with an impairment in insulin action, insulin tolerance testing demonstrated marked insulin resistance among IKKβ-MGKO mice when compared to pair-fed controls (Figures 2J and 2K). We next assessed tissue-level insulin action by measuring pAkt/Akt ratios in liver, epididymal white adipose tissue (eWAT), and skeletal muscle of HFD-fed mice fasted overnight and then injected with insulin (5 U/kg, i.p.) 15 minutes before sacrifice. Surprisingly, insulin signaling in the livers of IKKβ-MGKO mice was slightly improved compared to controls, whereas a similar analysis in eWAT showed no difference between genotypes (Figures 2L and 2M). In contrast, skeletal muscle insulin sensitivity was markedly reduced in IKKβ-MGKO mice compared to controls (Figures 2L and 2M). Together, these data support the hypothesis that increased hepatic gluconeogenesis may account for the elevation of fasting glucose (Figures 2H, 2I, and S1) while impaired muscle insulin signaling may explain the reduced insulin tolerance in IKKβ-MGKO mice (Figures 2J and 2K).

The hypothalamus contains glucose-sensing neurons that modulate the insulin sensitivity of peripheral organs via the sympathetic nervous system (Ruud et al., 2017; Steinbusch et al., 2015). Based on the impairment of insulin action observed in IKKβ-MGKO mice described above, we hypothesized that hypothalamic glucoregulatory mechanisms are altered by loss of microglial inflammatory signaling during DIO. Hypothalamic expression of genes implicated in neuronal glucose sensing (Dunn-Meynell et al., 2002; Roncero et al., 2004; Stanley et al., 2013; Toda et al., 2016) including glucose transporters 1 and 2 (GLUT1/*Slc2a1* and GLUT2/*Slc2a2*), *Gck*, and uncoupling protein 2 (*Ucp2*) were compared between IKKβ-MGKO mice and pair-fed controls after 16 weeks of HFD (Figure 3A). In the IKKβ-MGKO mice, unchanged levels of *Slc2a1* and elevated levels of *Slc2a2* suggested intact or even enhanced hypothalamic glucose uptake while reduced levels of *Gck* and *Ucp2* highlighted a potential functional reduction in hypothalamic glucose sensing (Figure 3A). To address this latter possibility, we quantified hypothalamic neuronal activation in response to a systemic glucose challenge in fasted IKKβ-MGKO mice and controls (2 g/kg i.p.; 60 minutes). IKKβ-MGKO mice had a marked reduction in the number of glucose-stimulated c-Fos^+^ cells in both the dorsomedial (DMH) and lateral (LHA) regions and a blunting of the glucose response in the ventromedial (VMH) nucleus (Figures 3B and 3C). Together, these data suggest that diet-induced microglial activation maintains hypothalamic glucose sensing during HFD feeding to preserve glucose tolerance in the setting of advancing DIO.

**Figure 3.**
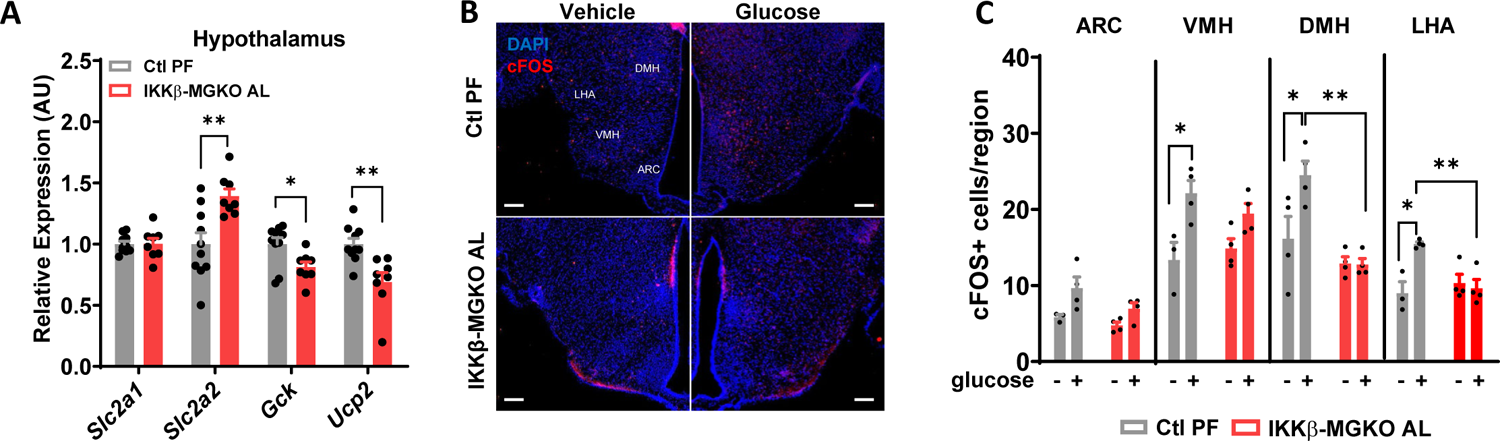
Hypothalamic glucoresponsiveness is impaired by loss of microglial IKKβ in obese mice. (A) Relative hypothalamic mRNA levels of glucose transporter 1 (*Slc2a1*) and 2 (*Slc2a2*), glucokinase (*Gck*), and uncoupling protein 2 (*Ucp2*) after 16 weeks of HFD in fasted IKKβ-MGKO AL normalized to levels of Ctl PF mice. N = 7/group. Student’s *t-*test, *p < 0.05, **p < 0.01. (B) Representative images of hypothalamic DAPI (blue) and c-Fos (red) immunostaining 1 hour after peripheral glucose administration (2 g/kg, i.p.) in 10 week HFD-fed mice. Arcuate nucleus, ARC; ventromedial hypothalamus, VMH; dorsomedial hypothalamus, DMH; lateral hypothalamic area, LHA. Scale bar = 100 μm. (C) Quantification of B. N = 3-4/group. Two-way ANOVA with Bonferroni post-hoc test, * p < 0.05, ** p < 0.01. All values are mean ± SEM.

### Increasing microglial inflammation improves glycemia in lean and obese mice

If silencing microglial IKKβ signaling worsens glucose tolerance during DIO, then enhancing microglial activation would be predicted to have the opposite effect. To test this hypothesis, we analyzed *Cx3cr1*^CreER/+^::*Tnfaip3*^fl/fl^ mice, a model of tamoxifen-inducible microglia-specific deletion of the NF-κB repressor protein A20 (Tavares et al., 2010; Valdearcos et al., 2017) (Figure 4A). We have previously reported that microglial A20 deletion causes immediate and spontaneous microglial inflammation, resulting in increased food intake and weight gain even on a CD (Valdearcos et al., 2017). For this study, the *Cx3cr1*^CreER/+^::*Tnfaip3*^fl/fl^ mice were first lethally irradiated with lead head shielding (to protect microglia and preserve the blood-brain barrier). The mice were subsequently transplanted with wild-type donor bone marrow in order to avoid confounding of metabolic studies by A20 deletion-mediated inflammation in peripheral monocytes, (A20-BMT-MGKO; Figure 4A). In the resulting chimeric mice, Cre induction deletes A20 from microglia but leaves the peripheral immune system unaffected (Valdearcos et al., 2017).

**Figure 4.**
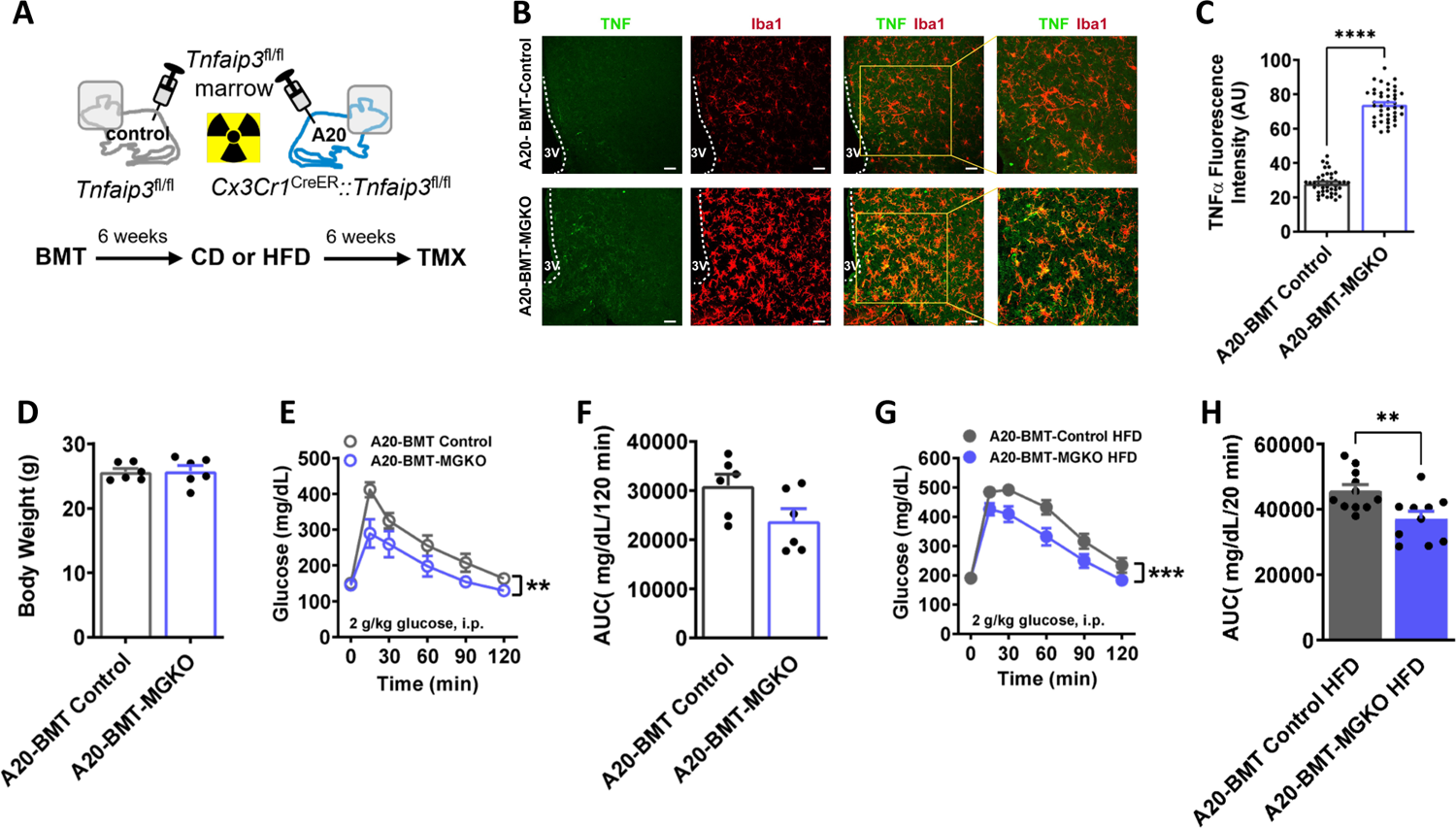
Derepression of microglial IKKβ/NF-κB signaling through A20 (*Tnfaip3*) deletion improves glucose tolerance. (A) Schematic representation of adoptive transfer of control *Tnfaip3*^fl/fl^ marrow into head-shielded control *Tnfaip3*^fl/fl^ (A20-BMT Control) and *Cx3Cr1*^CreER^::*Tnfaip3*^fl/fl^ (A20-BMT-MGKO) mice, followed by dietary intervention (CD or HFD) and inducible knockout (TMX = tamoxifen). (B) Hypothalamic TNF and Iba1 immunostaining in coronal sections from CD-fed BMT mice 1 week after TMX. Scale bars = 100 μm. (C) Quantification of B. Student’s *t*-test, **** p < 0.0001. (D) Body weight of CD-fed mice 1 week after TMX administration. N=6/group. (E) GTT (2 g/kg glucose, i.p.) and F) AUC 1 week after TMX in CD-fed mice. Two-way ANOVA, ** p < 0.01. (G) GTT (2 g/kg glucose, i.p.) and H) AUC in 6 week HFD-fed mice. Two-way ANOVA and Student’s *t*-test, ** p < 0.01, *** p < 0.001. All values are mean ± SEM.

1 week after tamoxifen treatment, CD-fed A20-BMT-MGKO mice displayed hypertrophic microglia in the mediobasal hypothalamus (Figure 4B) along with increased TNFα expression (Figures 4B and 4C), indicating a heightened state of inflammatory activation. Remarkably, A20-BMT-MGKO mice had improved glucose tolerance compared to BMT controls despite equal weights at this timepoint (Figures 4D-4F). In a separate cohort of mice fed HFD for 6 weeks, A20-BMT-MGKO mice had reduced obesity-associated glucose intolerance compared with control littermates (Figures 4G and 4H). Overall, these results reveal an unexpected effect of microglial inflammation to improve glucose tolerance both under physiologic conditions and during DIO.

### Diet-independent models of acute microglial inhibition and activation

Our results indicated an apparent dissociation between the detrimental impact of microglial activation on body weight regulation and its salutary effect on glucose homeostasis. To further explore this unexpected dichotomy, we sought a model of acute microglial silencing to avoid potential compensatory adaptations to permanent gene deletions. As a first approach, we administered the third generation tetracycline derivative minocycline, a widely used inhibitor of microglial inflammation (Banerjee et al., 2021; Garrido-Mesa et al., 2013; Yrjänheikki et al., 1998), into the CNS of 4 week HFD-fed rats. Rats receiving minocycline (10 μg i.c.v.) had an acute impairment in insulin tolerance compared with vehicle-treated controls (Figures S2A and S2B), analogous to the insulin resistant phenotype of the IKKβ-MGKO mice.

Next, we used the Designer Receptor Exclusively Activated by Designer Drugs (DREADD) chemogenetic system (Roth, 2016) to express the Gq-coupled hM3Dq (activating) receptor in microglia, which we hypothesized could provide a diet-independent means to trigger microglial activation *in vivo* using the synthetic DREADD agonist clozapine N-oxide (CNO). We first confirmed that *Cx3cr1*^CreER^::CAG-LSL-HA-Gq-DREADD-mCitrine mice (hM3D) treated with tamoxifen to activate Cre-mediated recombination had stable expression of the transgene in microglia up to 28 weeks later (Figure 5A). Moreover, expression of hM3D, as determined by qRT-PCR, was restricted to the CNS (Figure 5B; data not shown). As expected, hM3D mice and littermate controls (*Cx3cr1*^CreER/+^) showed equivalent weight curves on CD (data not shown) since the hM3Dq receptor remains inactive in the absence of CNO administration. To validate Gq-coupled signal transduction in this chemogenetic mouse model, we measured the change in intracellular calcium concentration ([Ca^2+^]_i_) in response to CNO treatment in primary microglia preloaded with the calcium indicator Fura Red. CNO treatment (0.01-0.1 μM) caused a rapid rise in [Ca^2+^]_i_, which remained elevated for several hours afterward (Figures 5C and 5D), consistent with prior reports using DREADDs in microglia (Binning et al., 2020; Grace et al., 2018; Saika et al., 2020, 2021; Yi et al., 2021). *In vivo*, CNO treatment (1 mg/kg; i.p.) increased microglial cell size but not cell number within the hypothalamus (Figures 5E-5G) and resulted in upregulation of cytokine and chemokine mRNA levels in FACS-isolated microglia (Figure 5H). Importantly, CNO treatment did not reduce 24-hour food intake or dark cycle ambulatory activity (Figures S3A-S3D), suggesting acute chemogenetic activation of microglia did not induce sickness behavior despite increasing inflammatory signaling (Figures 5E-5H).

**Figure 5.**
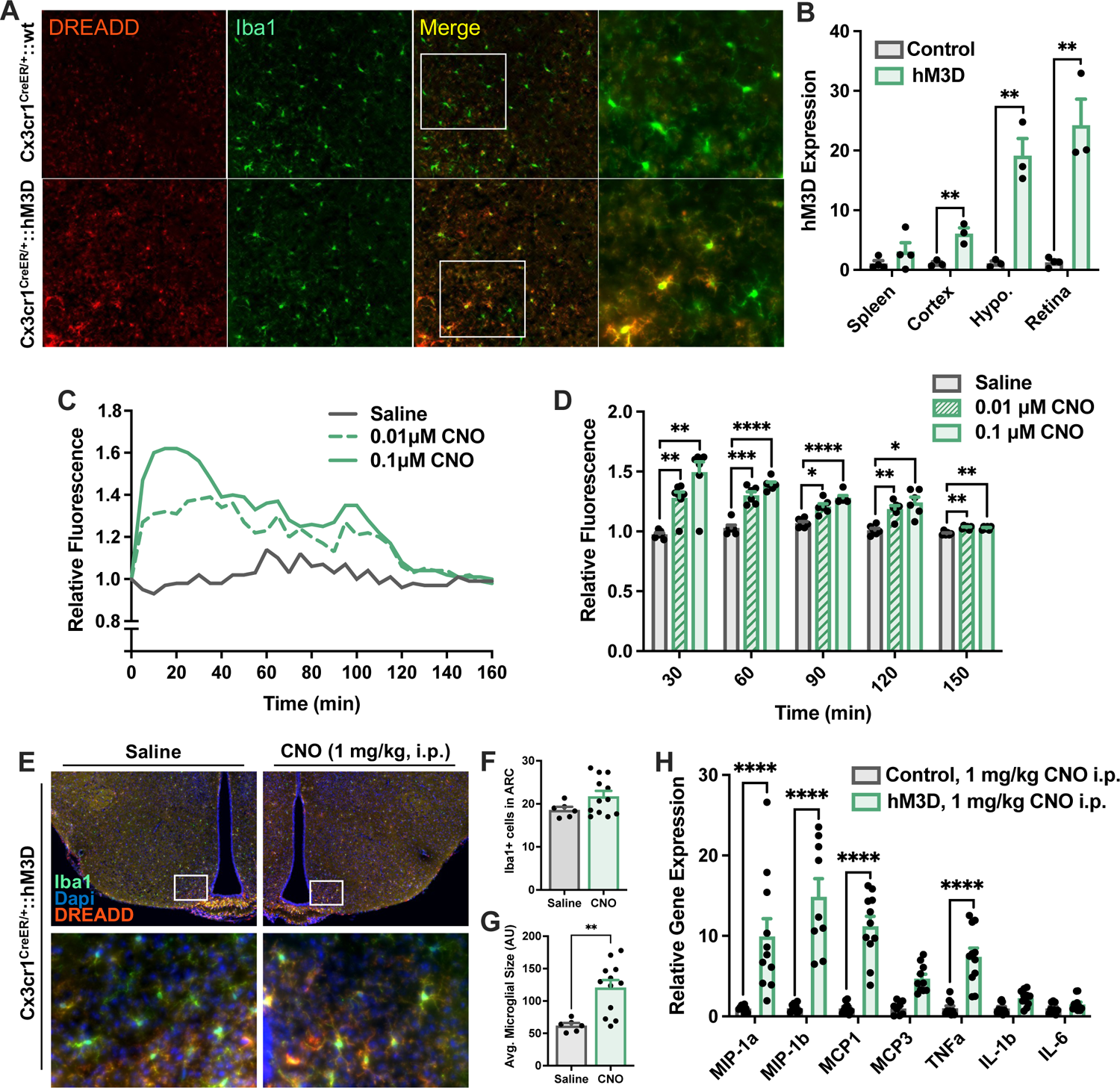
Chemogenetic activation via hM3D in microglia induces pro-inflammatory activation in mice. Cx3cr1^CreER/+^::LSL-HA-hM3Dq-citrine (hM3D) mice were generated as an inducible model of *in vivo* microglial activation. (A) Representative hypothalamic sections immunostained for the microglial marker (Iba1, green) and hM3D DREADD receptor (anti-HA tag, red) in tamoxifen-treated control and hM3D mice. (B) Quantification of hM3D transgene expression by qRT-PCR in CNS and peripheral tissues in hM3D mice normalized to control (Cx3cr1^CreER/+^) levels. N = 3-4/group. Student’s t-test, ** p < 0.01. (C) Fura Red was used to detect changes in intracellular calcium. Isolated hM3D primary microglia were treated with CNO (0.01 µM or 0.1 µM) or saline control over 150 minutes. Fura Red fluorescence of the microglial cells was measured using a plate reader and plotted normalized to 0 minute baseline. Mean traces shown for clarity. (D) 30 minute AUC bins of data in C. N = 6-7 replicates/treatment. Two-way ANOVA with Bonferroni post-hoc test, * p < 0.05, ** p < 0.01, *** p < 0.001, **** p < 0.0001. (E) Representative hypothalamic sections from hM3D mice immunostained for the microglial marker (Iba1, green), hM3D DREADD receptor (anti-HA tag, red), and DAPI nuclear stain in hM3D mice given CNO (1 mg/kg, i.p.) or saline and perfused 1 hour later. (F-G) Quantification of Iba1-positive microglia by (F) total number and (G) cellular size after CNO treatment. N = 6-12/group. Student’s t-test, **p<0.01. (H) Cytokine expression in FACS-isolated microglia from hM3D normalized to control levels in mice treated with CNO (1 mg/kg, i.p.). N = 9-12/genotype. Student’s *t*-test, **** p <0.0001. All values are mean ± SEM.

### Acute chemogenetic activation of microglia improves glucose tolerance in CD- and HFD-fed mice

Having validated the hM3D model, we next utilized systemic CNO administration to determine the effect of acute diet-independent microglial activation on glucose tolerance in CD-fed hM3D and control mice. While no differences were observed with vehicle treatment (data not shown), microglial activation with i.p. injection of 1 mg/kg CNO caused a significant but somewhat variable improvement in glucose tolerance in hM3D mice compared to controls (Figures 6A and 6B). At 5 mg/kg i.p., CNO treatment elicited a more consistent and robust glucose response, still without inducing sickness behavior (Figures 6C and 6D; data not shown). Prolonged treatment with CNO provided in the water (25 μg/ml; ∼75-100 μg daily) did not affect body weights (Figure 6E) but markedly improved glucose tolerance in hM3D mice compared with controls (Figures 6F and 6G). Interestingly, return of the animals to normal water reversed the benefit (Figures 6H and 6I), demonstrating that improved glucose tolerance depends on sustained microglial activation. Finally, to verify that the chemogenetic effect is mediated in the CNS, we compared different routes of CNO administration. Low-dose CNO treatment delivered centrally (1 µg i.c.v.; Figures 6J and 6K) but not systemically (1 µg i.p.; Figures S4A and S4B) improved glucose tolerance in CD-fed hM3D mice compared to controls, supporting a specific role for microglial activation in enhancing glucose disposal. This intervention also improved insulin sensitivity (Figures 6L and 6M), the parallel but opposite response to that obtained with minocycline treatment in rats (Figures S2A-S2C). These results demonstrate that acute induction of microglial inflammatory activation improves systemic glycemia independently of body weight or food intake alterations.

**Figure 6.**
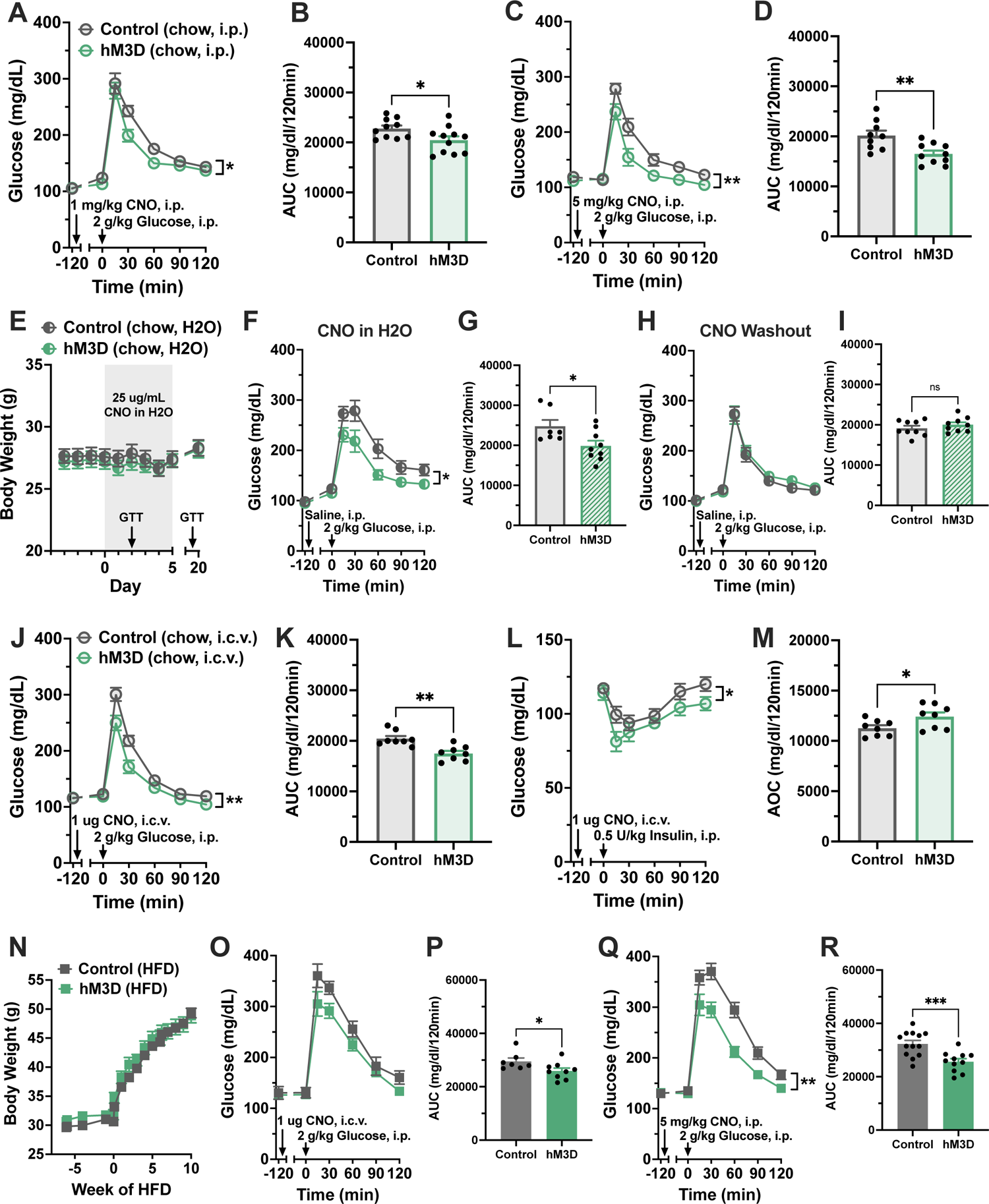
Acute pharmacogenetic activation of microglia improves glucose homeostasis in CD and HFD-fed mice. (A) Glucose tolerance test (GTT) and (B) AUC of CD-fed hM3D mice and littermate controls treated with systemic CNO (1 mg/kg, i.p., 2 hours prior; 2 g/kg glucose, i.p.), N = 11-14/group. (C) GTT and (D) AUC of CD-fed hM3D mice and controls treated with high-dose systemic CNO (5 mg/kg, i.p., 2 hours prior; 2 g/kg glucose, i.p.), N = 10-11/group. (E) Body weight of CD-fed hM3D mice and controls during chronic CNO administration via drinking water (25 μg/mL CNO and 1.92 μL/mL DMSO). Arrows indicate dates of GTTs performed in F and H. (F) GTT (2 g/kg glucose, i.p.) and (G) AUC following 2 days of CNO in the drinking water. A saline injection was given 2 hours prior to control for injection stress in other paradigms (10 μL/g, i.p.), N = 9/group. (H-I) Mice were maintained on CNO drinking water for 14 days then returned to normal water. (H) Follow-up GTT (2 g/kg glucose, i.p.) and (I) AUC were performed after 7 days of normal water, N = 9/group. (J-M) CD-fed hM3D mice and control littermates were injected with CNO (1 μg, i.c.v.) followed 2 hours later by metabolic testing. (J) GTT (2 g/kg glucose, i.p.) and (K) AUC, N = 8/group. (L) ITT (0.5 U/kg insulin, i.p.) and (M) area over the curve (AOC), N = 8/group. (N-R) hM3D mice and littermate controls were fed *ad libitum* HFD. (N) Body weights over 10 weeks of HFD. (O) GTT (2 g/kg glucose, i.p.) and (P) AUC for 4 weeks HFD-fed hM3D and control mice injected 2 hours prior with CNO (1 μg, i.c.v.), N = 8-10. (Q) GTT (2 g/kg glucose, i.p.) and (R) AUC for 8 weeks HFD-fed hM3D and control mice injected 2h prior with CNO (5 mg/kg, i.p.), N = 11-14. Two-way ANOVA and Student’s *t*-test, * p < 0.05, ** p < 0.01, *** p < 0.001. All values are mean ± SEM.

CD-fed mice have quiescent microglia at baseline, making them a logical model to assess the metabolic effects of microglial activation. However, they are also highly insulin sensitive and glucose tolerant, which constrains the degree to which microglial activation can further improve glycemic parameters. To overcome this limitation, we analyzed mice with established DIO, which are much more insulin resistant though already show a degree of microglial activation. As expected, hM3D mice fed a HFD gain weight at the same rate as control animals (Figure 6N) since the hM3D receptor remained inactive without CNO treatment. At 4 weeks of HFD feeding, acute central CNO administration (1 µg i.c.v.; Figures 6O and 6P) caused an improvement in glucose tolerance that was even more pronounced with high dose systemic treatment at 8 weeks of HFD (5 mg/kg i.p.; Figures 6Q and 6R). These data suggest that HFD-induced microglial activation can be further enhanced to offset obesity-associated glucose intolerance.

### Acute chemogenetic activation of microglia rescues glucose intolerance in obese mice through classical cytokine signaling

Microglia influence neuronal populations through multiple direct and indirect interactions, including the release of cytokines (Colonna and Butovsky, 2017; Prinz et al., 2019, 2021; Salter and Stevens, 2017). Given the findings in the IKKβ-MGKO and A20-BMT-MGKO mice (Figures 1-4) and the upregulated inflammatory gene expression in microglia from hM3D mice injected with CNO (Figure 5H), we hypothesized that chemokines or cytokines regulated by NF-κB— namely TNFα, IL-1β, IL-6, MIP 1α and 1β—might mediate the glucoregulatory effects of microglial activation. Notably, all of these molecules except TNF have been previously shown to improve systemic glycemia through signaling in the CNS (Chou et al., 2016; del Rey et al., 2006, 2016; Timper et al., 2017). However, neither central administration of an IL-6 blocking antibody (IL-6ab; 100 ng i.c.v.; Figures S5A and S5B), nor an IL-1β inhibitor (IL-1RA; 2 μg i.c.v.; Figures S5C and S5D), nor an antagonist to the MIP 1α/β receptor CCR5 (CCR5ant (maraviroc); 500 ng i.c.v.; Figures S5E and S5F) prevented the improvement of glucose tolerance in CNO-treated hM3D mice. In contrast, inhibition of TNFα signaling through central administration of etanercept (TNFnab (sTNFRII-Fc fusion protein); 2 μg i.c.v.) completely reversed the glycemic benefits of CNO-mediated microglial activation (Figures 7A-7C). Correspondingly, central infusion of low-dose TNFα (2 pmol i.c.v.) increased glucose disposal in CD-fed mice (Figures 7D and 7E). Together, these data implicate TNFα as a key mediator linking microglial activation to the regulation of glucose homeostasis.

**Figure 7.**
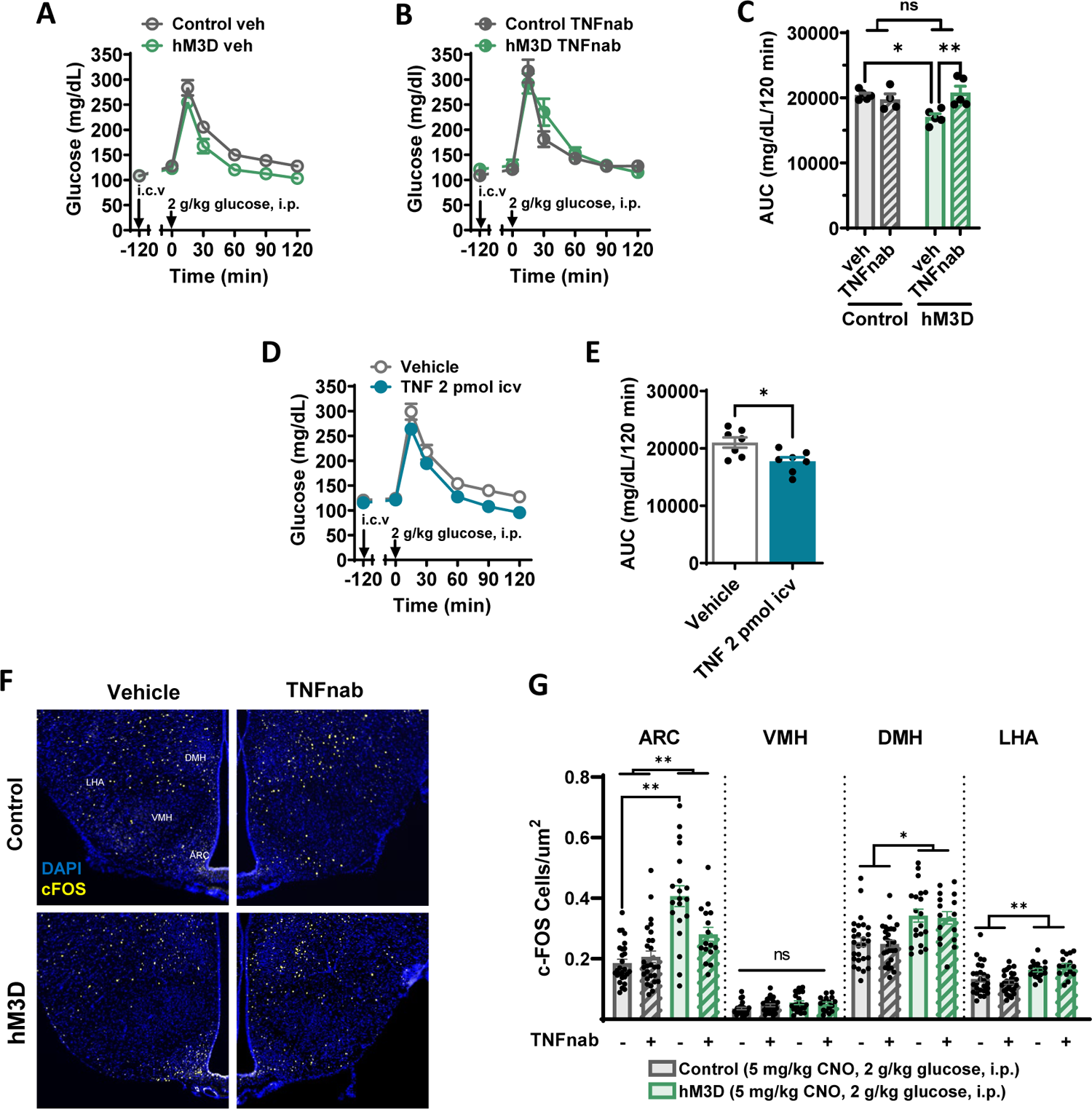
Microglial regulation of glucose homeostasis involves central TNF action. (A-B) GTTs (2 g/kg glucose, i.p.) performed in CD-fed control and hM3D mice 2 hours after simultaneous administration of CNO (1 mg/kg, i.p.) and (A) vehicle (IgG-Fc; 2 μg, i.c.v.) or (B) TNF neutralizing antibody etanercept (TNFnab) (2 μg, i.c.v.). Coinjection of i.p. CNO and i.c.v. veh/TNFnab is indicated by the arrow labeled “i.c.v.” N = 5-7/group. (C) AUC quantification of A and B. (D) GTT and (E) AUC in wild-type cannulated mice administered vehicle or TNFα (2 pmol, i.c.v.), N = 8/group. Two-way ANOVA with Šidák correction for multiple comparisons and Student’s *t*-test, * p < 0.05, ** p < 0.01. (F-G) CD-fed control and hM3D mice injected with CNO (5 mg/kg, i.p.) followed at 1h intervals by saline vehicle or TNFnab (2 μg, i.c.v.), glucose (2 g/kg, i.p.), and perfusion for IHC analysis. (F) Representative hypothalamic images of DAPI (blue) and c-Fos (yellow) immunostaining 1 hour after peripheral glucose administration in TNFnab and vehicle treated hM3D-activated mice. Arcuate nucleus, ARC; ventromedial hypothalamus, VMH; dorsomedial hypothalamus, DMH; lateral hypothalamic area, LHA. (G) Quantification of F. N = 5-8 mice/group, 1-6 sections/mouse. Results were analyzed in a mixed model controlling for repeated effects of individuals with a Tukey HSD post-hoc test. All values are mean ± SEM.

Finally, we sought to link the TNFα-mediated improvement in glucose tolerance in the hM3D model to the finding of reduced hypothalamic glucose sensing with loss of microglial inflammatory signaling in the IKKβ-MGKO mouse (Figure 3). Specifically, we hypothesized that induction of microglial activation in the hM3D model increases the activity of hypothalamic glucoresponsive neurons via TNF action. To address this possibility, we examined hypothalamic c-Fos staining in hM3D and control mice 60 minutes after serial treatment with CNO to activate microglia (5 mg/kg, i.p.) followed by vehicle or etanercept to block TNF action (TNFnab; 2 μg i.c.v.; 60 minutes after CNO) and glucose to mark glucoresponsive neurons (2 g/kg, i.p.; 60 minutes after i.c.v.). Compared with control mice, hM3D mice showed increased numbers of glucose-stimulated c-Fos^+^ neurons in both the DMH and LHA and an especially robust activation of neurons in the ARC (Figures 7F and 7G). Furthermore, central etanercept (TNFnab) administration significantly attenuated the ARC response (Figures 7F and 7G), demonstrating the involvement of TNF signaling in the enhancement of glucose sensing by microglial activation. Overall, these data support a model that posits microglial inflammatory modulation of hypothalamic glucose sensing as a mechanism to preserve glucose tolerance in the setting of advancing DIO.

## Discussion

Microglia provide key modulatory input in obesity pathogenesis, but an independent role in glucose homeostasis has not been previously established. Here we show with cell-specific mouse models that microglial activation caused by HFD feeding or chemogenetic induction drives an unexpected dissociation of energy metabolism and glucose tolerance. Despite significant protection against body weight gain, mice lacking microglial inflammatory signaling (IKKβ-MGKO) become more glucose intolerant than control mice during HFD feeding. Indeed, glucose tolerance and insulin sensitivity, along with indicators of neuronal glucose sensing, are actually *reduced* in IKKβ-MGKO mice relative to controls when the groups are weight matched. Conversely, microglial-specific deletion of A20 increases microglial inflammation and weight gain yet *improves* glucose tolerance in mice fed either a CD or a HFD. In a microglia-specific DREADD model, hM3D, that avoids the confounding effects of body weight differences or developmental compensation, microglial activation acutely improves glucose tolerance in both CD- and HFD-fed mice through a TNFα-dependent mechanism that increases hypothalamic glucose sensing. Together, these data comprise the first evidence that microglial activation offsets obesity-associated glucose intolerance despite contributing to DIO susceptibility.

Despite decades of research, the link between cellular inflammation and glycemia is still controversial. Contextual features lie at the core of this argument, namely the localization and exact nature of the inflammation. Fat-rich diets and obesity induce NF-κB transcriptional activity and the secretion of pro-inflammatory mediators in liver, WAT, and leukocytes (Saltiel and Olefsky, 2017). Earlier murine studies showed a clear connection between IKKβ-mediated NF-κB signaling in hepatocytes and diet-induced liver insulin resistance (Arkan et al., 2005; Cai et al., 2005). Further, DIO mice with myeloid-specific deletion of IKKβ maintain whole body insulin sensitivity (Arkan et al., 2005), implicating bone marrow-derived cells as disruptors of euglycemia. In contrast to these studies, transgenic mice with constitutively active IKKβ in cells expressing aP2—primarily adipocytes and macrophages—are protected from glucose intolerance (Jiao et al., 2012) while deleting IKKβ in adipocytes has a diabetogenic effect (Park et al., 2016). Moreover, local inactivation of the NF-κB transcriptional target TNFα in adipose tissue results in severe glucose intolerance and WAT remodeling, reminiscent of lipodystrophy (Wernstedt Asterholm et al., 2014). Additionally, Liu et al. demonstrated that virally-mediated liver IKKβ gain-of-function improves glucose homeostasis in adult *ob/ob* and DIO mice (Liu et al., 2016). Within the CNS, inflammatory signaling through IL-1β, IL-6, and MIP1/CCR5 has been implicated in improving glycemic parameters in lean and obese mice (Chou et al., 2016; del Rey et al., 2006, 2016; Timper et al., 2017). Combined with our current findings, this highlights the ongoing complexity of the inflammation-glucose tolerance debate and the need to develop a unified model for understanding the role of cellular, tissue-specific, and systemic versus CNS inflammation in diabetes pathogenesis.

Reactive glia are observed not only during the development of obesity but also in settings of altered glycemic regulation. Radiological analyses in human subjects show that gliosis correlates with insulin resistance independently of BMI (Berkseth et al., 2018; Schur et al., 2015; Thaler et al., 2012). Similarly, hyperglycemia in rats with poorly controlled diabetes is associated with gliosis in multiple brain regions (García-Cáceres et al., 2008; Nagayach et al., 2014). Surprisingly, insulin-induced hypoglycemia can also trigger microglial activation (Winkler et al., 2019). Together, these results could indicate that glucose levels regulate microglial activation rather than vice versa. However, live two-photon imaging indicates that microglia retain normal surveillance and phagocytic capacity even under aglycemic conditions (by using glutamate as an alternative fuel) (Bernier et al., 2020). Moreover, microglial silencing during hypoglycemia increases blood glucose (Winkler et al., 2019), similar to our findings of worsening glycemia in DIO rats treated with minocycline and the HFD-fed IKKβ-MGKO mice. These findings favor the hypothesis that microglia are key modulators of systemic glucose homeostasis, helping to maintain euglycemia in the face of environmental perturbations.

The central control of glucose homeostasis involves multiple brain regions and an ever expanding list of effector cells (Elizondo-Vega et al., 2015; Garcia-Caceres et al., 2016; Myers et al., 2021; Ruud et al., 2017; Steinbusch et al., 2015). Though our study highlighted a link between changes in microglial activation state, the activity of hypothalamic glucose sensing neurons, and systemic glucose homeostasis, these results do not preclude the involvement of other non-neuronal cell types or of neurons in extrahypothalamic regions in the microglial regulation of glucose homeostasis. The network of glucoregulatory neurons extends throughout the midbrain and hindbrain (Ter Horst et al., 2018; Jansen et al., 1997; Myers et al., 2021; Steinbusch et al., 2015), and our genetic and pharmacologic interventions affect microglia throughout the CNS. Unfortunately, there are few strategies to selectively target microglial gene expression in a regionally precise manner (Dumas et al., 2021; Maes et al., 2019). In addition, astrocytes participate in systemic glucose homeostasis regulation as well as brain glucose uptake and sensing (Garcia-Caceres et al., 2016) and have an inflammatory state that is intimately linked to microglial activity (Liddelow et al., 2017; Rothhammer et al., 2018). Thus, future work is needed to unravel the complex interactions between different glia cell populations such as astrocytes and microglia and determine how their input to neurons in the hypothalamus and other brain regions enables the fine tuning of systemic glucose homeostasis.

The hM3Dq-DREADD receptor activates Gq signaling upon administration of CNO leading to a rise in intracellular calcium and—in the case of neurons—an increased frequency of action potentials (Roth, 2016). Though largely applied to studies of neuronal functions, it has also been used to activate a variety of other cell types, both in the CNS and the periphery (Wang et al., 2021). While we and others have used the hM3Dq-DREADD in microglia as a means of cellular stimulation (Binning et al., 2020; Grace et al., 2018; Saika et al., 2020, 2021; Yi et al., 2021), it is a relatively unrefined mechanism of activation and lacks the nuances of environmental adaptation now known to be the cornerstone of microglial function in health and disease (Colonna and Butovsky, 2017; Prinz et al., 2019, 2021; Salter and Stevens, 2017). In this case, use of hM3D mice enabled us to selectively induce microglial activation and cytokine production, recapitulating key aspects of our A20 model and suggesting that microglial inflammatory mediators acutely impact glucose tolerance. The phenotypes observed with hM3D-mediated microglial activation generally paralleled the genetic models, but some differences in the findings may be due to the nature of the interventions. Chemogenetic activation in the hM3D model is acute and therefore its beneficial effect on glucose tolerance is likely mediated by changes in autonomic tone. In contrast, the A20 and IKKβ-MGKO models are genetic interventions that interface with diet exposure, feeding behavior, adiposity, and other elements of systemic metabolism to elicit tissue level changes that determine glycemia. Despite these differences, these two distinct approaches support the idea that microglial inflammatory signaling impacts a variety of glucoregulatory neurocircuits with far-ranging metabolic consequences. Future studies are needed to disentangle the similarities and differences between DREADD-mediated microglial activation and the complexity of microglial activation that occurs with HFD feeding and to determine whether microglial-targeted therapies are a viable avenue for glycemic interventions.

Acute hM3D activation with CNO increased microglial expression of inflammatory cytokines and chemokines, including MIP-1α/β and TNFα. Therefore, we tested a number of cytokines and chemokines downstream of the NF-κB signaling pathway to determine their involvement in the glycemic effects of chemogenetic microglial activaiton. Both IL6 and IL-1β have been shown to act within the hypothalamus to improve systemic glucose homeostasis (del Rey et al., 2006, 2016; Timper et al., 2017). In the context of acute microglial stimulation with DREADDs, neither IL-6 blocking antibody nor IL-1β inhibitor reversed hM3D-induced improvements in glucose tolerance. On the basis of our gene expression data, we then tested the effects of MIP-1α/β and TNFα inhibition. Hypothalamic CCR5 (the receptor for MIP-1 molecules) has been previously implicated in glucose regulation, with central blockade using a CCR5 antagonist impairing both central and peripheral insulin sensitivity and worsening peripheral glucose tolerance (Chou et al., 2016). Surprisingly, however, the CCR5 antagonist maraviroc did not limit the effects of microglial DREADD activation despite the expected worsening of glucose tolerance in control animals. Therefore, the impact of microglial activation on peripheral glucose homeostasis is apparently independent of hypothalamic MIP/CCR5 mechanisms involved in peripheral insulin signaling and glucose tolerance.

Instead of IL1, IL6, or MIP1, we identified TNFα signaling as a putative mediator for microglial regulation of glucose tolerance. This was unexpected, as TNFα secretion within the WAT of obese rodents and humans is considered a significant contributor to insulin resistance (Hotamisligil et al., 1993, 1995; Saltiel and Olefsky, 2017). Nevertheless, anti-TNFα therapy has paradoxically shown limited efficacy in improving glycemic parameters in obese individuals (Dominguez et al., 2005; Paquot et al., 2000). Could lack of effectiveness be attributable to an opposing function in the CNS? Intriguingly, recent work by Yi *et al*. shows that microglial activation and hypothalamic TNFα increase in lean mice during the fed state (Yi et al., 2017). The authors further demonstrate that TNFα acutely enhances evoked firing frequency in POMC neurons (Yi et al., 2017). Because POMC activity enhances glucose disposal and suppresses hepatic glucose production (Hill et al., 2010), this neuronal population is a potential mediator through which microglia-driven inflammation improves glucose homeostasis. This may arise from direct secretion of TNFα from microglia or perhaps another cell type, such as astrocytes, which have low TNFα expression in the resting state (Zhang et al., 2014) but can act downstream of microglia to amplify neuroinflammation (Liddelow et al., 2017). Despite our findings that pharmacological blockade of TNFα signaling reversed the improvement in glucose homeostasis of the hM3D model, it is likely that other microglial mediators also contribute. Future studies are needed to identify additional signaling molecules and determine the neuronal and glial targets involved.

There are a number of possible mechanisms by which CNS microglia may influence peripheral glucose homeostasis. Based on our c-Fos studies, microglial inflammatory signaling may potentiate neuronal glucose sensing in the hypothalamus. Alternatively, microglial activation may stimulate other neuronal effectors, altering downstream autonomic tone and thereby altering systemic glucose homeostasis. Neurons in the ARC are well-known to affect hepatic glucose production (HGP), particularly POMC neurons, which suppress HGP and increase peripheral glucose uptake and are responsive to TNFα (Hill et al., 2010; Pozo and Claret, 2018; Yi et al., 2017). In support of this hypothesis, our IKKβ-MGKO mice showed increased hepatic expression of gluconeogenic enzymes, suggesting an impairment in HGP suppression. We also found that IKKβ-MGKO mice had impaired insulin signaling in skeletal muscle. The PVN and other hypothalamic nuclei have been shown to affect skeletal muscle glucose uptake and disposal in a melanocortin signaling-dependent manner, although these regulatory circuits are still being enumerated (Pozo and Claret, 2018). Our models suggest that microglial inflammatory signaling may affect a number of these pathways, contributing to an overall phenotype of improved glucose homeostasis in response to microglial activation and worsened glucose tolerance when microglial inflammatory signaling is suppressed. These findings spotlight an important and novel aspect of physiology that should be explored in future studies and considered in the context of the parallel obesity and T2DM epidemics.

Here we demonstrate that microglia, the resident CNS innate immune cells, regulate energy balance and systemic glucose homeostasis in a reciprocal fashion. Loss of IKKβ in microglia reduces body weight but worsens glucose tolerance and exacerbates insulin resistance during a HFD challenge. Using multiple *in vivo* approaches, we also show that microglial activation enhances glucose disposal in both lean and obese animals. Overall, these findings identify a new central node in glycemic control and support the hypothesis that some aspects of microglial pro-inflammatory activation due to fattening diets may be metabolically beneficial. As a more complete understanding of this mechanism develops, strategies targeting microglial activation may serve as useful adjuncts to treatments for T2DM and other glucose-related disorders.

## ACKNOWLEDGMENTS

This work was supported by the American Diabetes Association (Pathway Award 1-14-ACE-51 to JPT), the NIDDK (K08 DK088872 and R01 DK119754 to JPT; R01 DK103175-02 to SKK; T32 DK007247 to JDD, KMN, and AWJ; and F32 DK108473 to JDD), and a UCSF Diabetes Family Fund Award to MV. In addition, services and support were provided by the Nutrition Obesity Research Center (DK035816) and Diabetes Research Center (DK017047) at the University of Washington.

## AUTHOR CONTRIBUTIONS

JDD, MV, and KMN designed and conducted experiments, analyzed data, prepared figures, and drafted the manuscript. AWJ, MDD, JF, AN, JB, RDF, and MR assisted in conducting experiments and provided input. SKK and JPT designed and supervised the studies and manuscript preparation.

## SUPPLEMENTAL FIGURE LEGENDS

### EXPERIMENTAL PROCEDURES

#### Rodents

Adult male Wistar rats were purchased from Charles River Laboratory (USA). All mouse studies used adult males from strains backcrossed onto the C57BL/6 background. Cx3cr1^CreER/+^ (strain #021160) and CAG-LSL-HA-Gq-DREADD-mCitrine (strain #026220) mice were purchased from The Jackson Laboratory. Conditional IKKβ deleter mice (*Ikbkb*^fl/fl^) were obtained from the laboratory of Dr. Michael Karin (Arkan et al., 2005). Conditional A20 deleter mice (*Tnfaip3*^fl/fl^) mice were obtained from Dr. Averil Ma (Tavares et al., 2010). Mice used for the study were as follows: 1) Cx3cr1^CreER/+^:: *Ikbkb*^fl/fl^ (IKKβ-MGKO); *Ikbkb*^fl/fl^ (Ctl) 2) Cx3cr1^CreER/+^::*Tnfaip3*^fl/fl^ (A20-MGKO); *Tnfaip3*^fl/fl^ (Control) 3) Cx3cr1^CreER/+^::CAG-LSL-HA-Gq-DREADD-mCitrine^+/-^ (hM3D); Cx3cr1^CreER/+^ (Ctl). Genotyping was performed by PCR using ear genomic DNA and the following primers: *Ikbkb* floxed allele (forward-CCT TGT CCT ATA GAA GCA CAA C; reverse-GTC ATT TCC ACA GCC CTG TGA); CreER allele (common forward-AAG ACT CAC GTG GAC CTG CT; mutant reverse-CGG TTA TTC AAC TTG CAC CA; wild type reverse-AGG ATG TTG ACT TCC GAG TTG); hM3D allele (mutant forward: CGC CAC CAT GTA CCC ATA C; mutant reverse: GTG GTA CCG TCT GGA GAG GA).

Unless otherwise specified, all mice were group-housed and age-matched with *ad libitum* access to water and diet in a pathogen- and temperature-controlled room with a 12:12h light:dark cycle (changed by the University of Washington facility to 14:10h light:dark during the hM3D studies). Mice were fed either a standard low-fat CD (LabDiet 5053) or a semi-purified HFD (60% fat (kcal), D12451, Research Diets Inc.). Anesthesia for euthanasia was by isoflurane or 100 mg/kg ketamine and 10 mg/kg xylazine. All procedures were performed in accordance with NIH Guidelines for Care and Use of Animals and were approved by the Institutional Animal Care and Use Committees at the University of California, San Francisco and the University of Washington.

### Induction of CreER-mediated recombination

Cre-ER-mediated recombination in the IKKβ-MGKO model was induced using 2 injections of tamoxifen 48-h apart (4 mg per dose in purified corn oil, s.c., Sigma). For the A20-BMT-MGKO mouse, recombination was elicited using 3 consecutive days of oral tamoxifen (5 mg per dose dissolved in purified corn oil, by gavage). For the hM3D model, a more prolonged tamoxifen protocol with 4 consecutive daily injections of tamoxifen (2 mg per dose in purified corn oil, i.p.) was used to increase recombination efficiency. For all models, control mice received the same tamoxifen regimen described above to ensure equivalent exposure. Except for the A20-BMT control and A20-BMT-MGKO mice, experiments were initiated at least 4 weeks after tamoxifen to allow for turnover and replacement of CX3CR1-expressing peripheral immune cells by newly born nonrecombined monocytes derived from the bone marrow. Microglia are long-lived and replenished through clonal proliferation so they retain the recombined allele indefinitely (data not shown; (Dumas et al., 2021; Goldmann et al., 2013; Parkhurst et al., 2013; Valdearcos et al., 2017; Yona et al., 2013)).

### Bone marrow transplantation

Bone marrow chimeras were created as described previously (Valdearcos et al., 2017). Briefly, 8 week-old A20-BMT control or A20-MG-BMTKO mice were anesthetized and individually placed in lead tubes (RPI) to shield their heads and necks from irradiation. The mice were lethally irradiated in two 5Gy doses given 3h apart and underwent bone marrow transplantation the next day by tail-vein injection with 3 x 10^6^ femoral and tibial bone marrow cells flushed from A20 control donor mice. Mice were allowed to recover for 6 weeks, followed by dietary intervention and tamoxifen induction of CreER-mediated recombination.

### Dietary Interventions

Adult male Wistar rats were given *ad libitum* HFD for the minocycline study. For studies using the *Ikbkb*^fl/fl^ mouse strain, two separate feeding paradigms were performed. For the strictly *ad libitum* experiment, 6 week-old *Ikbkb*^fl/fl^ (Ctl) and *Cx3cr1*^CreER/+^::*Ikbkb*^fl/fl^ (IKKβ-MGKO) mice were injected with tamoxifen (see above) and then switched to HFD or maintained on CD 4 weeks later at 10 weeks of age, with weekly body weight and food intake measurements. Glucose tolerance was assessed at week 10 of HFD. For pair feeding experiments, 10 week-old Ctl and IKKβ-MGKO mice were maintained as follows: group averages of 24-h intake of the *ad libitum*-fed IKKβ-MGKO mice were measured on the day prior to the pair-fed Ctl mice receiving 70% of their total daily food at 17:00 and the 30% at 09:00 the following day (Tschöp et al., 2011). All mice in the pair feeding study were handled twice daily with body weights and food intake recorded once daily. To avoid confounding by different timing of food intake between pair-fed and *ad libitum* groups, all mice were allowed free access to food for 2 days prior to glucose and insulin assessments. For A20 studies, 6 weeks after bone marrow transplantation (see details above), adult *Tnfaip3*^fl/fl^ (A20-BMT Control) and *Cx3cr1*^CreER/+^::*Tnfaip3*^fl/fl^ (A20-BMT-MGKO) mice were maintained on CD or switched to HFD for 6 weeks. Mice were then administered tamoxifen (see above) and subjected to measures of glucose homeostasis 1 week after. For hM3D studies, cohorts of male Cx3cr1CreER/+::LSL-HA-hM3Dq-Citrine (hM3D) and littermate control Cx3cr1CreER/+ mice underwent tamoxifen administration at 6 to 8 weeks old (see above). 2 weeks later, one cohort underwent lateral ventricular cannulation (see method below) while other cohorts were left uncannulated. All hM3D mice on CD diet underwent measures of glucose homeostasis at least 4w after tamoxifen. For DIO studies, several CD cohorts were switched to HFD feeding at ∼16-20w of age and assessed beginning 4w after diet start.

### Chemogenetic studies of glucose homeostasis

Cohorts of CD and HFD-fed hM3D and Ctl mice received training i.p. injections and a baseline GTT (see method below) using i.p. saline vehicle prior to clozapine N-oxide (CNO) studies. Stock CNO was prepared in DMSO and diluted to 1% DMSO in saline to final concentrations of 0.03 mg/mL, 0.1 mg/mL, or 0.5 mg/mL; mice received 10 μL/g body weight yielding 0.3 mg/kg, 1 mg/kg, or 5 mg/kg doses as stated. On experimental days, food was removed by 9am, followed 2h later by administration of saline or CNO (0.3-5 mg/kg, i.p.; Hello Bio or Tocris). GTT, ITT, pyruvate tolerance, or glucose-stimulated insulin secretion was performed 2h after CNO administration (see below for methods). Animals were allowed at least 1 week recovery between studies.

For CNO water studies, we adapted a published protocol (Ewbank et al., 2020). A CD-fed cohort of male hM3D mice and Ctls was single-housed at 12 week-old and trained on water bottles for 2 weeks. Baseline daily water and food intake measures were taken for 1 week and then all mice were switched to CNO in the water (25 μg/ml; dissolved at 13 mg/ml in DMSO then diluted in sterile water to final conc of 0.2% DMSO). CNO water stored at 4°C was distributed fresh to animal bottles every 1-3 days and shielded from light throughout the experiment. Mice were maintained on CNO water for 14 days with GTT using i.p. saline vehicle performed on day 2, then were returned to normal water with repeat GTT using i.p. saline vehicle on day 7 of normal water.

### Intracerebroventricular Administration

Adult male Wistar rats were given *ad libitum* HFD for 4 weeks prior to i.c.v. cannulation. 6 week-old male hM3D and Ctl mice were given tamoxifen and underwent cannula implantation at 8 weeks of age. For the TNF study, a separate cohort of 8 week-old male C57BL/6 mice underwent cannula implantation. Steel guide cannulas (Alzet, DURECT Corp.) were implanted into the lateral ventricle of mice and third ventricle of rats using standard coordinates (Dorfman et al., 2017; Thaler et al., 2009). Animals were allowed to recover for 2 weeks and received multiple i.c.v. training injections prior to experiments. A 10 μL Hamilton syringe and sterile flexible tubing was used to inject conscious unrestrained rodents. Body weight and food intake was monitored daily throughout the experiments and animals were excluded if adverse reactions to injections occurred. For the rat studies, 3 doses of minocycline (10 μg, 2 μL total per injection) were administered: twice on the day prior and once on the day of the ITT (final dose 1h before insulin administration). For mouse i.c.v. CNO studies, all injectates were 2 μL of volume and performed 2h prior to GTT or ITT. For CNO alone studies, hM3D and Ctl mice received single injections of saline vehicle or CNO (1 μg; Tocris). For cytokine inhibition studies, hM3D and Ctl mice received CNO (1 mg/kg, i.p.) at the same time as either vehicle (recombinant human IgG-Fc, 110-HG-100, R&D Systems) or neutralizing TNFα antibody (2 μg; sTNFRII-Fc (etanercept), Amgen); vehicle (goat IgG, AB-108-C, R&D Systems) or goat anti-mouse IL-6 antibody (250 ng; AF-406, R&D); vehicle (0.1% BSA in sterile PBS) or recombinant human IL-1RA (100 μg; 200-01RA, Peprotech); vehicle (1% DMSO in saline) or CCR5 antagonist (500 ng; maraviroc, Tocris). For the TNF study, wild-type mice received i.c.v. vehicle (1% BSA in sterile PBS) or recombinant mouse TNF peptide (2 pmol; 5178, Cell Signaling) administered 2h prior to GTT. Peptide and inhibitor doses used were determined according to effective ranges from published studies (Nilsberth et al., 2009). For quantification of glucose stimulated c-Fos in hM3D mice, i.c.v. injection of vehicle (sterile saline) or neutralizing TNFα antibody (2 μg, 1 μL total; sTNFRII-Fc (etanercept), Amgen) was performed two hours prior to sacrifice (between i.p. injections of CNO 5 mg/kg and glucose 2 g/kg).

### Glucose Tolerance Testing

Glucose tolerance tests (GTT) were performed as described previously (Douglass et al., 2017b) with some modifications. Animals were fasted for 4 h (09:00-13:00) or overnight (21:00-09:00, pair-feeding experiments) prior to glucose administration (D-(+)-glucose, 30% solution, 2 g/kg, i.p.). Glucose was measured in tail blood by hand-held glucometer (FreeStyle Lite). Total area-under-curve (AUC) was calculated by the trapezoid rule. Glucose-stimulated insulin secretion was performed as in the GTT with minor modifications. Glucose (2 g/kg; by gavage) was administered followed by collection of tail blood at 0 and 15 minutes for insulin ELISA.

### Insulin Tolerance Test

Animals were fasted for 4h (09:00-13:00) or overnight (21:00-09:00, pair-feeding experiments) and administered insulin (Humulin, Lilly) at specified doses (0.5-1.5 units per body weight; i.p.). Glucose was measured in tail blood by glucometer. Area-over-curve (AOC) glucose levels were calculated by the trapezoid rule.

### Pyruvate Tolerance Test

Animals were fasted 6 hours (09:00-15:00) and administered pyruvate (2 g/kg, i.p., Sigma). Glucose was measured in tail blood by glucometer.

### Plasma Analyses

Plasma insulin was measured by ELISA in mice and rats (mouse and rat insulin ultra-sensitive assay, Crystal Chem).

### Energy Balance and Body Composition

To measure short term food intake in IKKβ-MGKO AL and Ctl PF mice, food intake was determined 4h into the dark cycle on at least 3 separate occasions during the pair feeding regimen. For food intake studies in hM3D mice, CD-fed animals had food removed 4h prior to the start of the dark cycle and were injected with saline or CNO 30 minutes prior to dark cycle onset and food return. Food and body weight were determined at 0 and 24h. 3 days later, mice were crossed over to the opposite treatment and the measurements repeated.

Body composition analysis including fat mass and lean mass measurements was performed at the NIDDK-funded Nutrition Obesity Research Center (NORC) Energy Balance Core at the University of Washington using EchoMRI in awake, conscious mice. For the pair feeding study, mice were sacrificed after 14 weeks of HFD with immediate weighing of gonadal fat pads.

### Ambulatory activity

Ambulatory activity was assessed with assistance from the NORC Energy Balance Core. To quantify the effect of chemogenetic microglial activation on activity, adult male hM3D mice underwent implantation of transponders (Starr Life Science Corp, Oakmont, PA) in the peritoneal cavity. Briefly, mice were anesthetized using isofluorane and the abdominal region was shaved. After sterilization with 70 % EtOH and betadine solution, a 3/4-inch incision was made in the skin and peritoneal membrane, and the transponder inserted into the peritoneal cavity. The wound was closed with sutures. The animals were treated with Buprenex and allowed 1 week recovery before single housing in a reverse light-cycle room (12:12 hour light/dark cycle starting at 01:00). After 1 week acclimation to handling and i.p. injections, mice were injected with either saline or CNO (1 mg/kg; i.p.) 2h after dark cycle onset. Transponder signals enabled activity counts to be sensed every minute by a receiver positioned underneath each cage and analyzed using VitalView software. For graphical presentation, activity traces were smoothed by plotting averages of 10 minutes per data point. Binned activity counts were computed as the mean ± SEM of all values obtained throughout each 30 minutes period post-injection.

### Microglial isolation

Microglia were isolated from whole brains of adult male control and HM3D mice. Two hours after an i.p. injection of CNO (1 mg/kg), mice were anaesthetized and transcardially perfused with ice cold PBS. The brains were removed and minced in cold media containing Accutase and DNase I. After incubation on ice for 45 minutes, the mixture was passed through a 45 μm filter to obtain a single cell suspension, mixed with a 40% Percoll (GE Healthcare) suspension, and the myelin was removed after pelleting the cells with centrifugation at 400 x *g.* The cell suspension was incubated with an anti-CD11b-FITC antibody (Cell Signaling Technologies) for 30 minutes on ice and immediately isolated by fluorescence activated cell sorting (FACS) for the live (DAPI^lo^) myeloid (CD11b^+^) population. Sorted cells were collected in RNA lysis buffer and subsequently processed for qRT-PCR.

### Primary microglia culture

Primary cultures of cortical mixed glia from P0-2 mouse pups were prepared as previously described (Douglass et al., 2017b). After 14 days in culture media (high glucose MEM, 10% fetal bovine serum, Glutamax, mCSF, penicillin/streptomycin) flasks were shaken at 220 rpm for 6 hours at 37°C to release microglia. The cells were pelleted, washed with Dulbecco’s PBS (DPBS) and plated onto 96-well plates. To induce recombination, all wells were treated for 48 hours with 5 μM 4-hydroxytamoxifen (4-OHT, Sigma), following by two DPBS washes and 4-OHT free culture media. Cells were serum starved overnight prior to treatment with serum free media with or without LPS (100 ng/ml; Sigma) for 6 hours. Cells were then washed twice with DPBS and lysed for RNA extraction and subsequent analyses.

### hM3D microglial calcium signaling

4-hydroxytamoxifen treated primary microglia from hM3D mice were shaken at 300 rpm for 3 hours (37 °C), plated and left to rest for 5-6 hours in a 96 well plate (2 x 10^4^ cells/well). Cells were washed with warmed 1x PBS and media was replaced overnight with MEM (Gibco #51200-038) supplemented with 320 µM glucose, 1x GlutaMAX (Gibco #35050061) and 1x penicillin-streptomycin (Gibco #10378-016). Cells were loaded in the dark for 1h at 37 °C with 30 µM Fura Red/AM (Invitrogen #F3021) made up in 0.33 % DMSO (Sigma #D1436) and 0.065 % Pluronic F-127 (Sigma #P2443). After Fura incubation, cells were washed with warmed PBS and treated with complete MEM supplemented with saline, 0.01 µM, or 0.1 µM CNO (Tocris) and taken immediately to the plate reader. The plate was incubated at 37 °C for 3.5 hours. Readings were taken every 5 minutes using an excitation of 485 nm and an emission of 675 nm. The average 675 nm reading taken from non-template wells (F_0_, wells incubated with Fura Red but without cells) was deducted from well readings (F_t_) and this value was divided by F_0_ (Wendt et al., 2015). Data smoothed using an average of five readings per 5 minute time point and expressed as a ratio of saline (F_t_-F_0_)/F_0_ at 0 hours (Wendt et al., 2015). Data was binned over 30 minutes and analyzed for significance compared to saline treated time-matched controls.

### Immunohistochemistry

Anesthetized mice were transcardially perfused with ice cold PBS and then 4% paraformaldehyde, followed by dissection of tissues which were then postfixed overnight (4°C). Brain, liver, pancreas, retina, and spleen were immersed in 30% sucrose for 48 hours, embedded in optimal cutting compound (OTC), and immediately frozen on dry ice and stored at −80°C. Floating sections were cut on a cryostat, blocked for 1 hour with 2.5-4% normal donkey serum in PBS containing 0.5% Triton X-100, and incubated with primary antibodies overnight at 4°C. Primary antibodies used were Iba1 (1:1000 rabbit polyclonal, Wako; 1:1000 goat polyclonal, Novus Biologicals NB100-1028), GFP (1:2000 chicken polyclonal, Aves Labs, Inc.), HA (1:1000, Cell Signaling Technology C29F4), TNF (1:300 mouse monoclonal, Abcam, ab1793), and c-Fos (1:4000 rabbit polyclonal, Oncogene 4191-1-1). Sections were then incubated for 2h at room temperature with the appropriate secondary antibodies (1:500 dilutions): donkey anti-rabbit AlexaFluor594 or AlexaFluor647, donkey anti-goat AlexaFluor 488, goat anti-mouse AlexaFluor 488, and goat anti-chicken AlexaFluor 488 (all from Invitrogen). Sections were briefly stained with DAPI (1:10,000, Invitrogen D1306) and brushed onto glass slides. Images were captured on an Eclipse E600 epifluorescence microscope equipped with a color digital camera (Nikon) or a confocal laser-scanning microscope (Leica TCS SP5).

The Histology and Imaging Core of the University of Washington Diabetes Research Center (DRC) performed the immunohistochemistry staining for insulin on pancreatic sections. All immunohistochemistry staining was performed on formalin-fixed paraffin embedded mouse pancreas tissues. First, the slides were baked for 30 minutes at 60°C and deparaffinized on the Leica Bond Automated Immunostainer (Leica Microsystems, Buffalo Grove, IL). All subsequent steps were performed at room temperature. Blocking consisted of Leica Peroxide Block for 5 minutes. Additional blocking occurred with 10% normal goat serum in TBS for 20 minutes. The anti-mouse insulin antibody (1:4000 guinea pig polyclonal; Dako, A0564) was diluted in Leica Primary Antibody Diluent and applied for 30 minutes. The secondary antibody (1:1500 unconjugated rabbit anti-guinea pig, Abcam, ab102356) was diluted in 5% NGS and TBS and incubated for 8 minutes. Next a horse radish peroxidase-conjugated tertiary antibody (goat polyclonal anti-rabbit IgG) was applied for 8 minutes. Antibody complexes were visualized using Leica Bond Mixed Refine (DAB, 3,3’-diaminobenzidine) detection 2X for 10 minutes. Tissues were counterstained with hematoxylin for 4 minutes followed by two rinses in water and dehydrated through graded alcohol to xylene. Slides were mounted with a synthetic mounting media and images were acquired with a Zeiss AxioImager brightfield microscope. Unless otherwise specified all reagents were obtained from Leica Microsystems.

### Histological Quantification

Iba1^+^ cells were counted manually from matched sections within prespecified regions of interest using ImageJ software. Microglial cell size was determined using a thresholding protocol (ImageJ) followed by densitometric quantification. TNF immunoreactivity (ir) was measured by converting images to binary and applying a minimum threshold to normalize non-specific staining, followed by analysis of mean pixel intensity in the specified region of interest. For IKKβ-MGKO glucose-responsive neuron quantification, c-Fos-positive cells were counted manually in defined hypothalamic nuclei from anatomically matched sections between −1.22mm to −2.30mm from bregma (Paxinos Mouse Brain Atlas). For quantification of glucose-stimulated c-Fos in hM3D mice, 5-7 sections/mouse were stained and imaged from −1.4 mm to −2.1 mm from bregma, and regions of interest (ROIs) were defined based on DAPI staining for hypothalamic nuclei of interest (ARC, VMH, DMH, and LHA). A binary mask of c-Fos images was created, thresholded, size filtered for minimum and maximum particle size, and analyzed as particles per μm^2^ per ROI (Fiji, ImageJ). Sections were scored for GFP fluorescence by a drug and genotype-blinded researcher to determine recombination efficiency with exclusion of hM3D sections with low GFP from the final analysis. Insulin immunoreactivity in pancreatic islets was measured by dividing the total pixel area positive for insulin by the hematoxylin-positive total tissue area.

### Western Blotting

Mice were injected i.p. with insulin (5 U/kg body weight) or saline and sacrificed by CO_2_ asphyxiation 15 minutes later. Liver, skeletal muscle (tibialis anterior), and epididymal fat tissues were collected and immediately frozen on dry ice. Phospho-Akt (Ser473) and total Akt were analyzed by standard immunoblotting procedures. Frozen samples were homogenized by probe sonication in ice-cold RIPA buffer containing protease and phosphatase inhibitors (Fisher). Protein concentration was determined by BCA assay (Fisher). Samples were loaded (20 μg protein per well) onto SDS-PAGE and electrophoresed for 1 hour at 120 volts. Proteins were transferred onto PVDF membranes for 1 hour at 120V using a BioRAD submerged electrophoresis apparatus. Membranes were blocked for 1 hour in 5% bovine serum albumin (BSA) and tris-buffered saline with 0.1% Tween-20 (TBST), incubated overnight with primary antibodies (pAkt Ser473 (Cell Signaling); total Akt (Cell signaling); beta actin (Cell Signaling) at 4°C with gentle rocking, and incubated at room temperature with secondary antibody (goat anti-rabbit conjugated HRP) for 2 hours. Membranes were treated with ECL reagent for 1 minute and visualized using the BioRAD ChemDoc digital imaging system.

### Real-Time Quantitative PCR

For RNA extraction, animals were anesthetized using CO_2_ and rapidly decapitated. Liver tissue was rapidly removed, and hypothalamic blocks were dissected, immediately followed by freezing on dry ice and storage at −80⁰C until RNA extraction. Total RNA was extracted using Trizol or RNeasy Mini kit according to manufacturer’s instructions (Qiagen) and reverse-transcribed with High Capacity cDNA Reverse Transcription Kit (Applied Biosystems). Transcript levels were measured by semiquantitative real-time PCR on an ABI Prism 7900 HT (Applied Biosystems) machine and SYBR green detection of amplicons. Data was analyzed using the sequence detection system software (SDS version 2.2; Applied Biosystems) and relative mRNA abundance was normalized to 18S. Primer sequences were designed using NCBI Primer3 as follows:

18S (internal control) forward: 5’-CGG ACA GGA TTG ACA GAT TG-3’,

18S (internal control) reverse: 5’-CAA ATC GCT CCA ACT AA-3’;

TNFα fwd: 5’-CAT CTT CTC AAA ACT CGA GTG ACA A-3’,

TNFα rev: 5’-TGG GAG TAG ATA AGG TAC AGC CC-3’;

Il1β fwd: 5’-TAC AAG GAG AGA CAA GCA ACG ACA-3’,

Il1β rev: 5’-GAT CCA CAC TCT CCA GCT GCA-3’;

Il6 fwd: 5’-GTG GCT AAG GAC CAA GAC CA-3’,

Il6 rev: 5’-GGT TTG CCG AGT AGA CCT CA-3’;

MIP-1α fwd: 5’-CTGCCCTTGCTGTTCTTCTC-3’

MIP-1α rev: 5’-GATGAATTGGCGTGGAATCT-3’

MIP-1β fwd: 5’-CTGCCCTCTCTCTCCTCTTG-3’

MIP-1β rev: 5’-AGGAAGTGGGAGGGTCAGAG-3’

MCP1 fwd: 5’-CTTCTGGGCCTGCTGTTCA-3’

MCP1 rev: 5’-CAAGCCTACTCATGGGCTTC-3’

MCP3 fwd: 5’-CCATGAGGATCTCTGCCACG-3’

MCP3 rev: 5’-ACCCACTTCTGATGGGCTTC-3’

Slc2a1 fwd: 5’-CCCCGTCCTGCTGCTATTG-3’

Slc2a1 rev: 5’-GCACCGTGAAGATGATGAAGAC-3’

Slc2a2 fwd: 5’-TGTTGGGGCCATCAACATGA-3’

Slc2a2 rev: 5’-AACATGCCAATCATCCCGGT-3’

Ucp2 fwd: 5’-AAGCAGTTCTACACCAAGGGC-3’

Ucp2 rev: 5’-TCTCGTGCAATGGTCTTGTAG-3’

Gck fwd: 5’-CAA GCT GCA CCC GAG CTT-3’

Gck rev: 5’-TGA TTC GAT GAA GGT GAT TTC G-3’

G6pc fwd: 5’-TCAACCTCGTCTTCAAGTGGATT-3’

G6pc rev: 5’-CTGCTTTATTATAGGCACGGAGCT-3’

Pck1 fwd: 5’-GGC GGA GCA TAT GCT GAT CC-3’,

Pck1 rev: 5’-CCA CAG GCA CTA GGG AAG GC-3’

Gq DREADD fwd: 5’-TCT GGC AAG TGG TCT TCA TC-3’

Gq DREADD rev: 5’-TTA CCA GGA TGT TGC CGA TG-3’

### Statistics

Analyses were performed using GraphPad Prism 6.0 and 9.3 and JMP Pro 16.0.0 software. Data are presented as mean ± SEM. Two-group comparisons used unpaired two-tail Student’s *t* test except for within subjects crossover experiments when paired Student’s *t* tests were used. For comparisons of more than two groups or for GTT/ITT analyses, two-way ANOVA or linear mixed models were used with repeated measures when appropriate, followed by Bonferroni, Šidák, or Tukey post-hoc testing. Glucose AUC and fat mass values were compared by linear regression. *P* values < 0.05 were considered significant.

---

**Figure S1.**
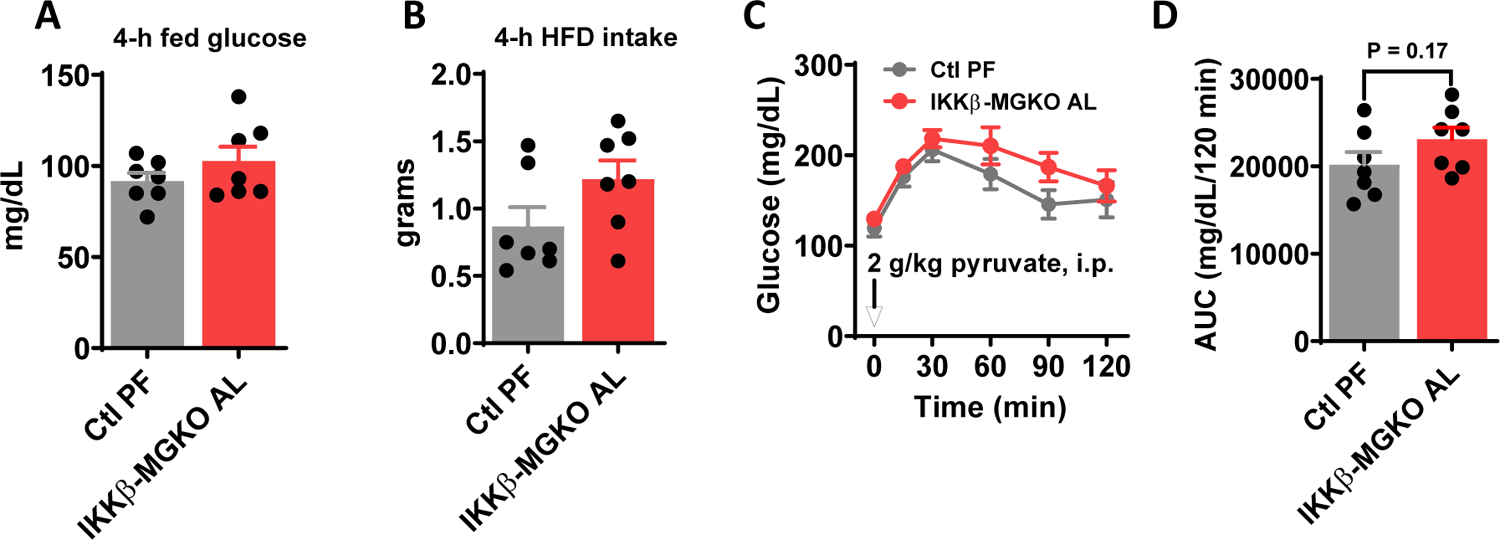
HFD pair-feeding does not impact glucose or food intake but deficiency of microglial inflammatory signaling tends to increase gluconeogenesis. (A) Blood glucose and (B) food intake in IKKβ-MGKO mice fed *ad libitum* (IKKβ-MGKO AL) vs. pair-fed control (Ctl PF). Measurements were obtained 4 hours into the dark cycle. N = 7/group. C) Pyruvate tolerance test (2 g/kg pyruvate, i.p.) and D) total glucose AUC at 11 weeks HFD in IKKβ-MGKO mice fed *ad libitum* vs. pair-fed control.

**Figure S2.**
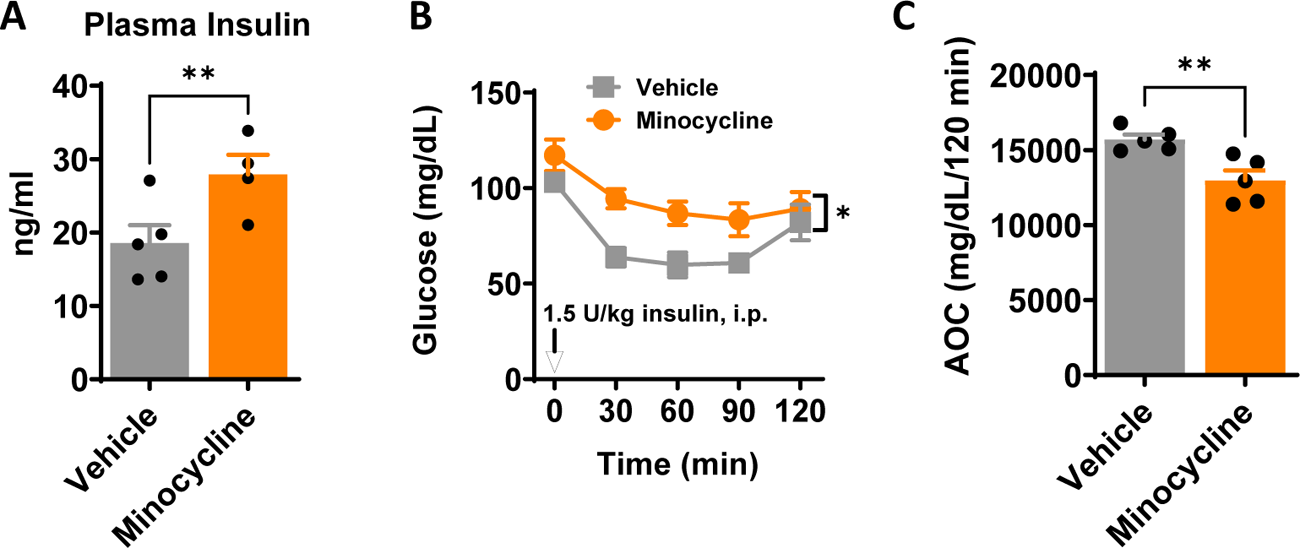
Pharmacological inactivation of microglia worsens insulin resistance during DIO in rats. 3V-cannulated male Wistar rats (N = 5/group) were fed HFD for 4 weeks and treated with minocycline (10 μg, i.c.v.) or saline vehicle administered 3 times over 2 days. (A) Plasma insulin levels at t = 0 of ITT. (B) ITT (1.5 U/kg insulin, i.p.) performed 1h after the third dose of minocycline or vehicle. (C) Total glucose AOC for the ITT. Values are mean ± SEM. 2-way ANOVA with Bonferroni post-hoc test and Student’s *t*-test, * p < 0.05, ** p < 0.01.

**Figure S3.**
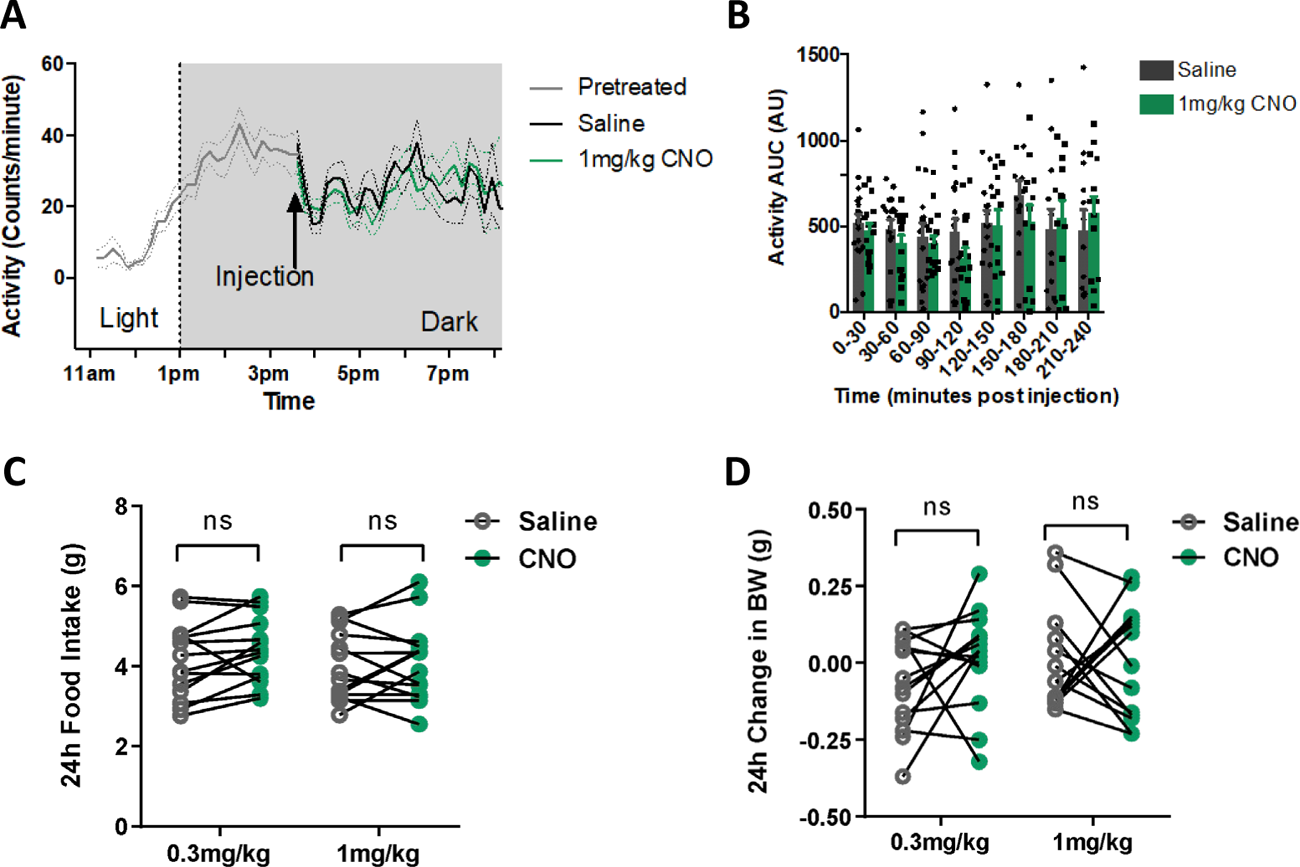
Acute CNO activation of microglial hM3D does not induce sickness behavior. (A) Dark cycle ambulatory activity was measured after saline or CNO injection (1 mg/kg, i.p.) into hM3D mice. Pretreatment line represents average of both groups of hM3D mice before injection (indicated by arrow). Dashed lines represent ± SEM. (B) 30 minutes AUC bins of activity for 240 minutes after injection. N = 13-19/group. (C) Food intake and (D) body weight were measured 24 hours after saline or CNO injection (0.3 mg/kg or 1 mg/kg, i.p.) using a within subjects crossover design. N = 14/group.

**Figure S4.**
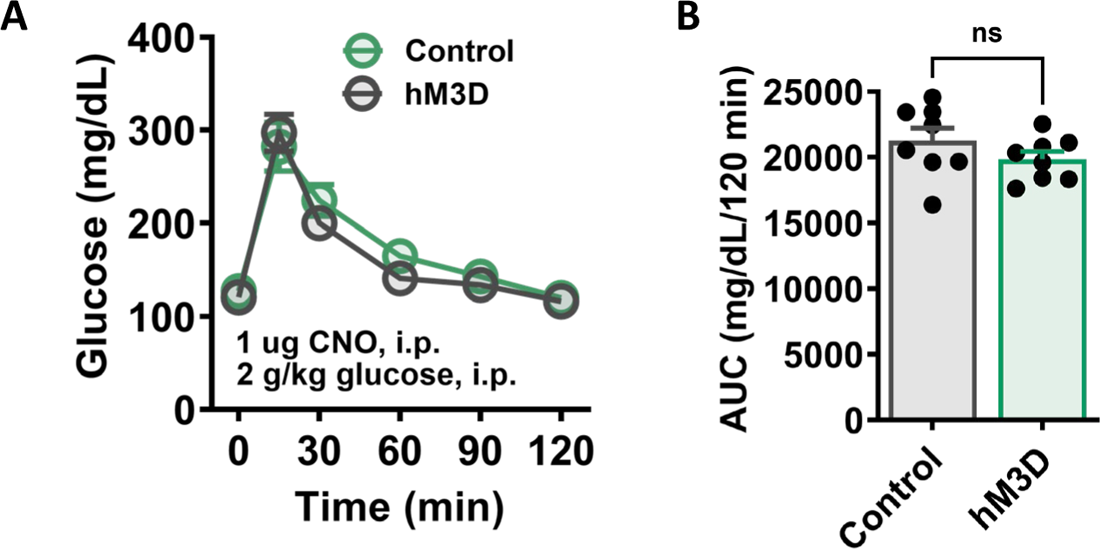
Subthreshold peripheral CNO administration does not improve glucose tolerance in HM3D mice. (A) GTT (2 g/kg glucose, i.p.) and (B) AUC in CD-fed hM3D and Ctl mice administered systemic CNO at the same dose used for i.c.v. experiments (1 μg, i.p., 2 hours prior). N = 8/group. All values are mean ± SEM.

**Figure S5.**
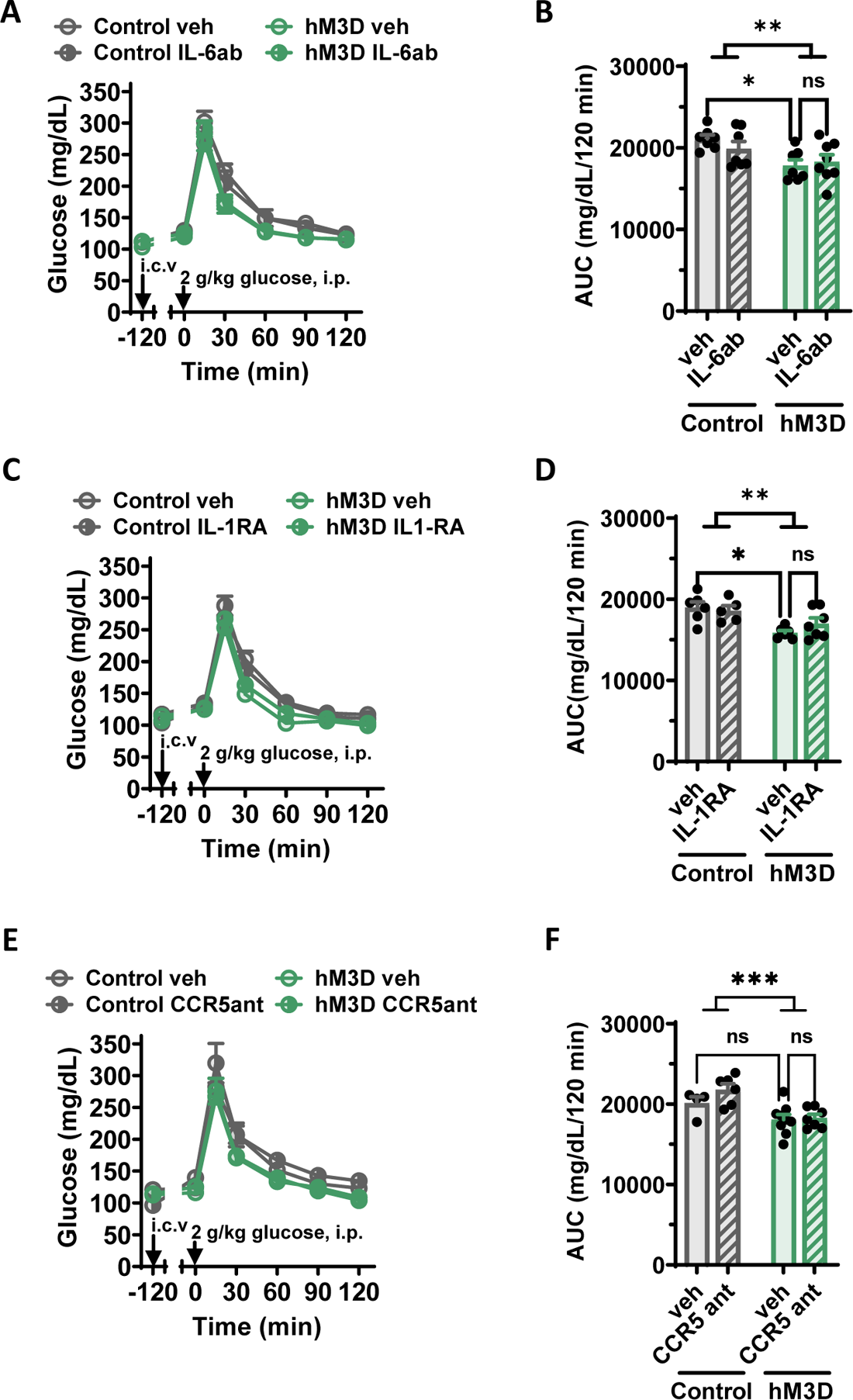
Central blockade of IL-6, IL-1, and CCR5 does not impact glucose tolerance during chemogenetic activation of microglia. (A-C) CD-fed control and hM3D mice with indwelling lateral ventricle cannulas were injected with CNO (1 mg/kg, i.p.) and i.c.v. treatment listed below (indicated in figures by arrow labeled i.c.v.) 2 hours prior to all GTTs (2 g/kg glucose, i.p.). (A) GTT and (B) AUCs of mice administered vehicle or IL-6ab (150 ng, i.c.v.). (C) GTT and (D) AUC of mice administered vehicle or IL-1RA (100 ng, i.c.v.). (E) GTT and (F) AUC of mice administered vehicle or CCR5 antagonist maraviroc (CCR5ant) (500 ng, i.c.v.). Two-way ANOVA with Šidák correction for multiple comparisons, * p < 0.05, ** p < 0.01, *** p < 0.001. All values are mean ± SEM.

